# A local translation program regulates centriole amplification in the airway epithelium

**DOI:** 10.1101/2022.01.18.476821

**Authors:** Helu Liu, Huijun Li, Zhihua Jiang, Shibo Jin, Rui Song, Ying Yang, Jun Li, Jingshu Huang, Xiaoqing Zhang, Xuesong Dong, Munemasa Mori, Marvin J Fritzler, Lin He, Wellington Cardoso, Jining Lu

## Abstract

Biogenesis of organelles requires targeting of a subset of proteins to specific subcellular domains by signal peptides or mechanisms controlling mRNA localization and local translation. How local distribution and translation of specific mRNAs for organelle biogenesis is achieved remains elusive and likely to be dependent on the cellular context. Here we identify Trinucleotide repeat containing-6a (*Tnrc6a)*, a component of the miRNA pathway, distinctively localized to apical granules of differentiating airway multiciliated cells (MCCs) adjacent to centrioles. In spite of being enriched in TNRC6A and the miRNA-binding protein AGO2, they lack enzymes for mRNA degradation. Instead, we found these apical granules enriched in components of the mRNA translation machinery and newly synthesized proteins suggesting that they are specific hubs for target mRNA localization and local translation in MCCs. Consistent with this, *Tnrc6a* loss of function prevented formation of these granules and led to a broad reduction, rather than stabilization of miRNA targets. These included downregulation of key genes involved in ciliogenesis and was associated with defective multicilia formation both in vivo and in primary airway epithelial cultures. Similar analysis of Tnrc6a disruption in yolk sac showed stabilization of miRNA targets, highlighting the potential diversity of these mechanisms across organs.

**Highlights:**

1. *Tnrc6a* is expressed in the lung selectively in differentiating multiciliated cells (MCC) adjacent to centrioles.
2. TNRC6A localizes to apical granules containing AGO2, miRNAs and their targets, but lacking mRNA degradation enzymes.
3. TNRC6A granules are enriched in components of the mRNA translation machinery and show evidence of concentrated newly-synthesized proteins
4. Loss of *Tnrc6a* in the lung leads to reduction, not stabilization of miRNA targets.
5. *Tnrc6a* is required for efficient centriole amplification and multicilia formation.

## Introduction

Biogenesis of subcellular organelles requires efficient delivery and concentration of subsets of proteins to specific intracellular sites for prompt assembly^1,2^. Targeting proteins to specific subcellular compartments with signal peptides or by translation of localized mRNAs is known to be involved in this process^3–5^. mRNA subcellular localization and local translation are evolutionary conserved and efficient mechanisms to create functional and structural asymmetries in cells by rapidly producing encoded proteins when and where required^3^. The prevalence and precise subcellular localization of certain mRNAs in association with specific organelles suggest a significant contribution of local translation^4^. Mechanisms generating asymmetric distribution of proteins in polarized cells are functionally relevant in regulating multiple aspects of cell behavior, including proliferation, migration and differentiation.

Centrioles are core structures of centrosomes involved in nucleation and formation of mitotic spindles critical for cell division^6^. Centrioles also function as basal bodies for the assembly of primary cilia and multicilia^6,7^. During multiciliogenesis, each differentiating multiciliated cell (MCC) generates hundreds of centrioles. A number of mRNAs localize and concentrate in centrioles or centrosomes^3,8,9^. Concentration of proteins and mRNAs in centrosomes might be achieved by targeting actively translating polysomes^8^. In addition, centriolar proteins are suggested to recruit the protein translation machinery that regulates the translation of specific mRNAs^10^. Still, little is known about the mechanisms of mRNA localization and local translation in centriole amplification. It is also unclear how microRNAs contribute to this process.

Trinucleotide repeat containing 6a (TNRC6A) is a direct partner of Argonaute proteins (AGOs), which play a critical role in miRNA-induced mRNA degradation^11^. AGOs and TNRC6A have been reported in randomly-distributed cytoplasmic granules known as P-bodies, which are enriched in translation suppressors and mRNA degradation enzymes, but lack protein synthesis machinery^12^. TNRC6A and AGO2 can be also detected in other cytoplasmic ribonucleoprotein granules, such as stress granules and neuronal transport granules^13–15^. Cell culture studies show *Tnrc6a* present in most cell types, however analyses of developing and adult tissues in vivo show that levels of *Tnrc6a* expression can differ markedly^16–18^.

In an effort to gain further insights into the tissue distribution and processes associated with *Tnrc6a* and miRNAs, here we identified*Tnrc6a* in the lung selectively expressed in MCCs of conducting airways. We found TNRC6A protein asymmetrically distributed in these cells in a not previously identified population of apical granules, which also contained AGO2. However these granules differed from P-bodies as they lacked mRNA degradation enzymes and instead were enriched or closely associated with components of the mRNA translation machinery and newly-synthesized proteins. Loss of function of *Tnrc6a* prevented these granules from forming and resulted in broad reduction of miRNA targets, including key components of the centriole biogenesis machinery required for multicilia formation. Our study suggests that this novel class of TNRC6A-containing apical granules can act as local translation hubs required to mediate large-scale production of centriolar proteins in MCCs. Our data also supports the idea that key components of the miRNA pathway can be utilized for mRNA recruitment and localization required for local translation.

## Results

### *Tnrc6a* is selectively expressed in multiciliated cells of the lungs

We have previously generated a *Tnrc6a*^gt/+^ reporter mouse carrying a gene trap insertion of β-galactosidase in the *Tnrc6a* locus to investigate expression and function of *Tnrc6a* in mouse development^18^. In the process of further characterizing these mutants, we performed marker analysis and X-gal staining in the adult lungs and found a striking selective pattern of β-galactosidase activity in the airway epithelium (**Fig. 1a)**. Little to no signals were present in the alveolar epithelium and overall in the mesenchymal compartment. This pattern was similarly observed for *Tnrc6a* as determined by in situ hybridization (**Fig. 1b)**. Moreover, β-galactosidase activity co-labeled with acetylated α-tubulin (Ac-α-tub), indicating that *Tnrc6a* was selectively expressed in MCCs of the airway epithelium (**Fig. 1c)**.

**Fig. 1.**
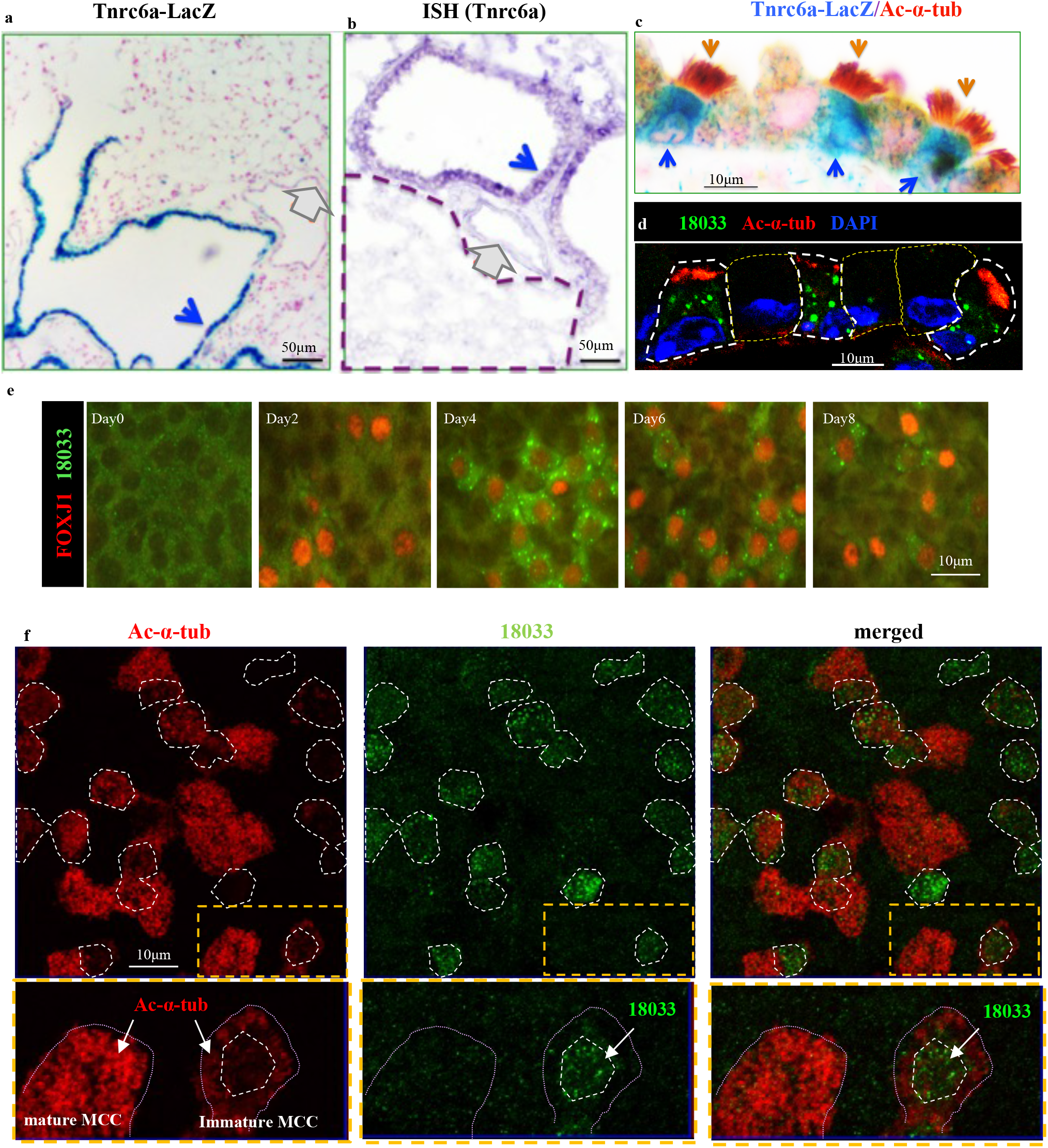
*Tnrc6a* is selectively expressed in multiciliated cells (MCCs) of airways in the mouse lungs. a) β-galactosidase staining of *Tnrc6a*^gt/+^ reporter mice showing selective expression in the airway epithelium (blue arrow), compared to vascular (gray arrow) and alveolar structures. b) *Tnrc6a* in situ hybridization confirming expression in airway epithelium (blue arrow), but not in vessels (red arrow) or alveoli (outlined by dashed line). c) Strong β-galactosidase signals (blue arrow) in MCCs labeled by Ac-α-tub (brown arrow) compared to other airway cell types. d) Immunofluorescence (IF) with human index serum (18033: green) detects TNRC6A in MCCs (Ac-α-tub: red). e) Time course of detection of TNRC6A-containing granules in MCCs (FOXJ1+) differentiating from adult airway progenitors in air-liquid interface (ALI) cultures. f) Distinct abundance of cytoplasmic TNRC6A expression (18033, dotted area and arrow) in immature and mature MCCs in ALI day4 cultures, as shown by strong or weak Ac-α-tub signals, respectively. Lower panels enlarged from boxed areas (yellow).

To further confirm expression and cell type specificity, we performed immunofluorescence (IF) staining of lung sections with the human index serum (termed as 18033), widely used for detection of TNRC6A protein^19,20^. This showed cytoplasmic signals restricted to Ac-α-tub positive cells, further corroborating TNRC6A selectivity to MCCs (**Fig. 1d)**. To investigate when these granules are established and their association to multiciliogenesis, we analyzed airway epithelial progenitors (basal cells) from adult mouse trachea undergoing differentiation in air-liquid interface (ALI) cultures^21^. Analysis of TNRC6A (18033) and FOXJ1, an early marker of MCC cell fate^21^ showed TNRC6A signals first detected at ALI day2 in emerging FOXJ1+ cells. By day3-4, as these cells underwent large-scale centriole amplification to initiate multiciliogenesis^21^, TNRC6A+ cytoplasmic granules became abundant in MCCs (**Fig. 1e**). Signals were subsequently diminished in more mature MCCs, which by then expressed strong Ac-α-tub **(Fig. 1f)**.

### TNRC6A is enriched in a novel class of apical epithelial granules that lack mRNA degradation enzymes

To gain additional insights into the subcellular distribution of these granules in MCCs, we analyzed a series of Z-stack images of ALI day 4 cultures double-stained with the human index serum and FOXJ1. This revealed an overall population of granules heterogeneous in size, randomly distributed throughout the cytoplasm of MCCs but also identified a population of apically-localized granules of smaller size in these cells. Immunostaining using a monoclonal TNRC6A antibody (4B6) and acetylated alpha tubulin showed these apical granules underneath the multicilia of differentiating MCCs (**Fig. 2a,b)**. Their localization in MCCs was further confirmed in ALI cultures transduced with a lentiviral vector expressing a TNRC6A-EGFP fusion protein^21^ (**Fig. 2c)**.

**Fig. 2.**
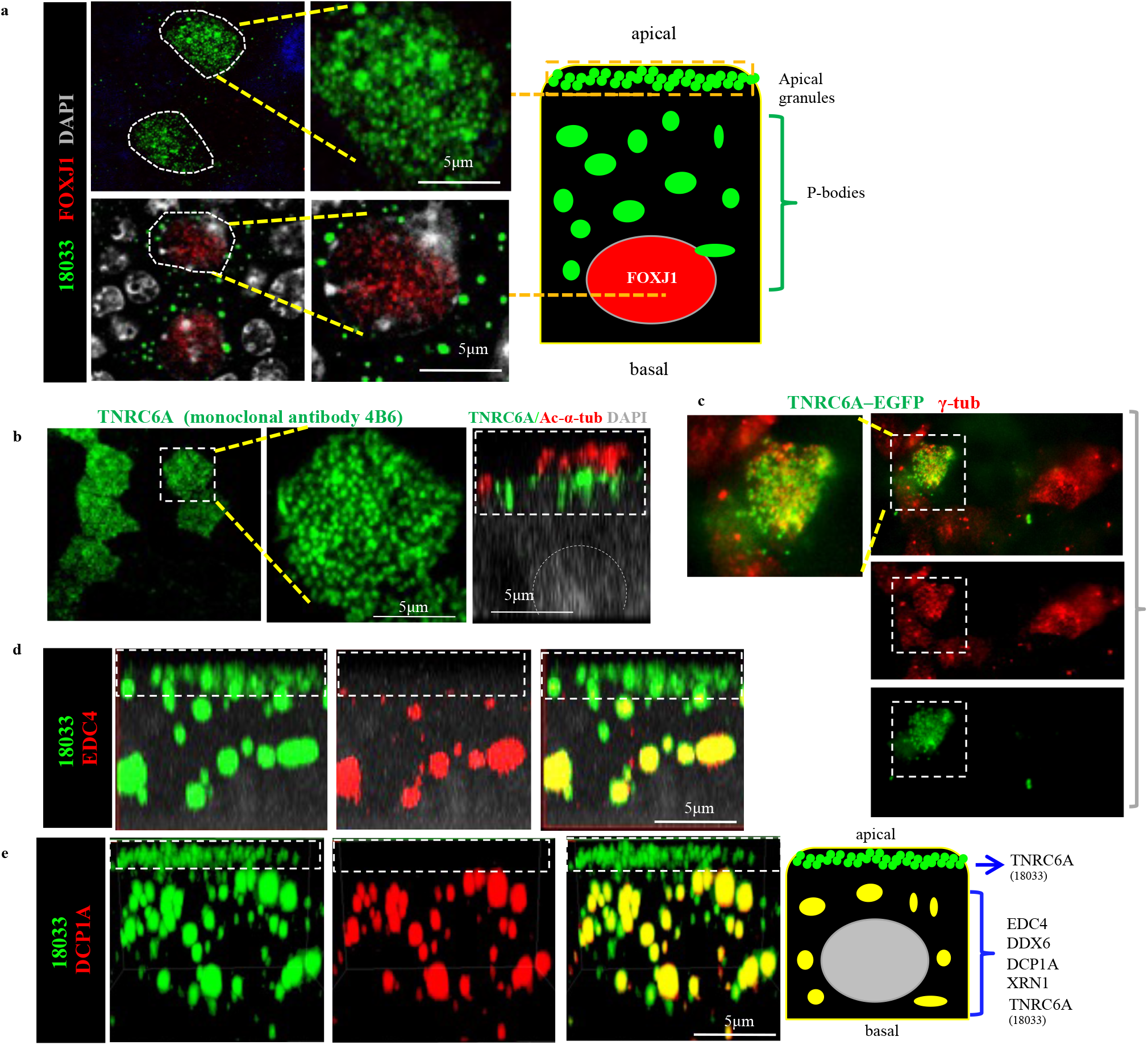
TNRC6A protein concentrates in apically-localized granules that lack mRNA degradation enzymes. a) Apical granules and randomly-localized granules (P-bodies) detected by human index serum (18033) in differentiating MCC. Dashed lines depict a plane of Z-stack image from the apical or basal areas of the same MCC (diagram). b) TNRC6A monoclonal antibody (clone#:4B6) identifies apical granules underneath the multicilia (Ac-α-tub). X-Z projection of confocal z-stack images. c) TNRC6A-EGFP fusion protein expressed by a lentiviral vector concentrates in apical granules of MCC identified by co-labeling with γ-tub (boxed area) in ALI day3 culture d-e) Co-staining of human index serum with EDC4 or DCP1A in a single MCC showing double-labeling largely in P-bodies instead of the apical granules (dashed box) Diagram depicts distinct components of these granules (See also Suppl. Fig.1)

The human index serum is also known to label P-bodies (processing bodies), cytoplasmic ribonucleoprotein (RNP) granules, which contain TNRC6A and various RNA pathway enzymes involved in decapping and degradation of mRNAs^22^. To further identify potential differences in the populations of MCC granules, we stained ALI cultures with human index serum and antibodies against components of the P-bodies. IF for 18033 double-labeled with EDC4, DCP1A, DDX6 and XRN1 showed these signals highly expressed in the randomly-localized population of cytoplasmic granules (P-bodies), but largely undetectable in the apical granules that we also identified using the 4B6 antibody in the same MCC (**Fig. 2b,d,e, Supplementary Fig.1)**. TNRC6A is a known partner of AGO2^12^. Co-IF staining of human index serum with AGO2 showed co-localization of AGO2 in both classes of granules in MCC (**Fig. 3a**). Notably, lentiviral gene transduction of AGO2-EGFP fusion protein showed the strong EGFP signals enriched apically underneath the cilia of differentiating MCCs (**Fig. 3b**).

**Fig. 3.**
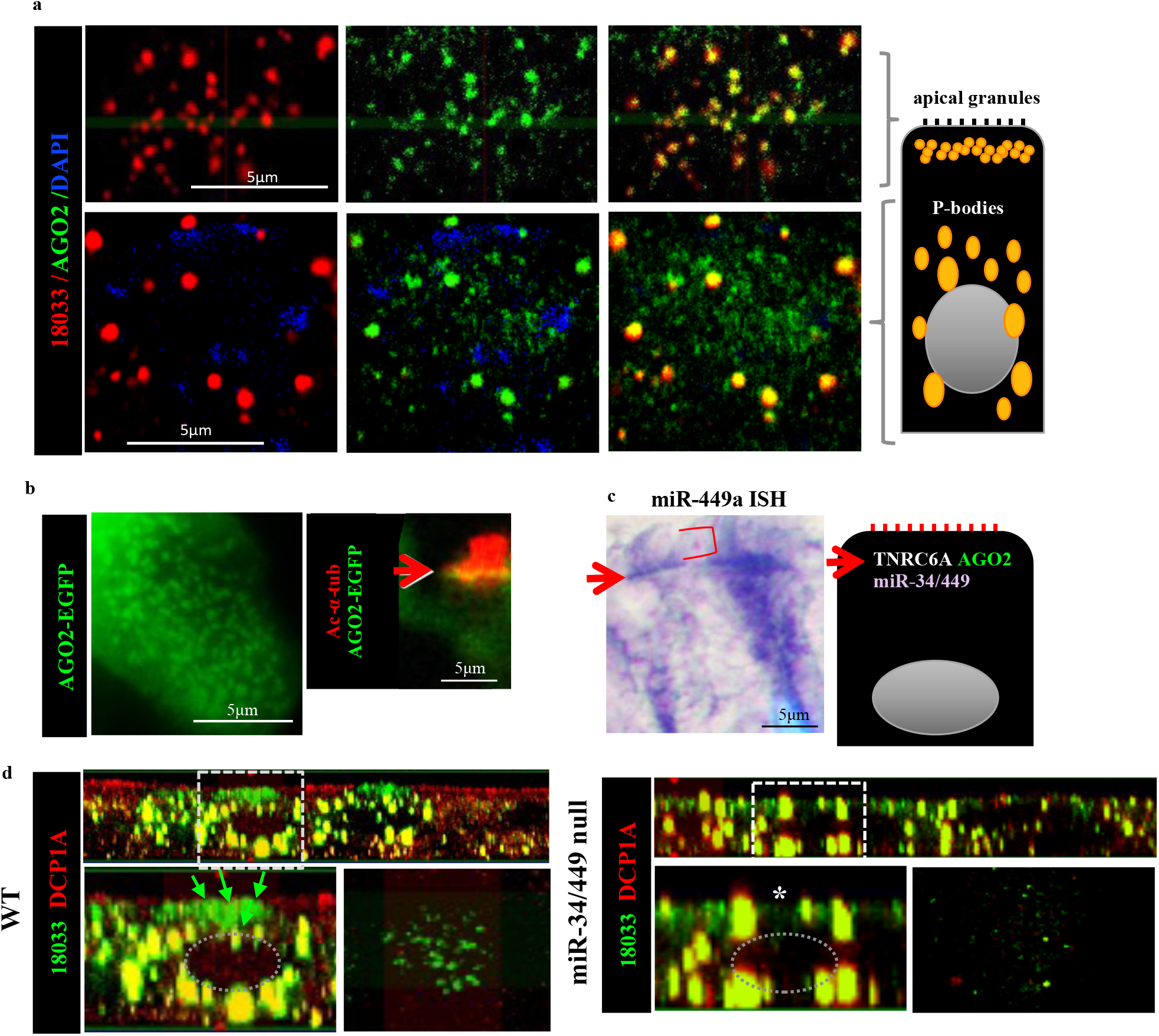
AGO2 and miRNAs are present in TNR6a-containing apical granules in differentiating MCCs. a) Co-staining of AGO2 (clone 11A9) and TNRC6A (18033, human index serum) showing double-labeling in apical granules and P-bodies within the same MCC. b) AGO2-EGFP fusion protein produced by a lentiviral vector in differentiating MCC in ALI cultures showing apical granules underneath the emerging cilia (red, arrow points to AGO2-EGFP fusion protein underneath Ac-α-tub-labeled cilia). c) In situ hybridization (ISH) showing miR-449a enriched apically (arrow) underneath the cilia (red bracket). d) Co-staining human index serum (green) and DCP1A (red) in ALI day 4 cultures from WT controls (left) and miR-34/449 triple null mice (right). Boxed area in top panels enlarged in bottom panel (YZ projection and a corresponding apical Z-stack image). WT: Abundant single-labeled TNR6A apical granules (arrows: 18033, green) and double-labeled larger granules (yellow 18033-DCP1A: P-bodies). miR-34/449 null ALI cultures: notable decrease in abundance of TNR6A apical granules (*, green) not seen in the P-body population (yellow double-labeled granules). Nuclei represented in grey (diagrams).

This supported the idea that TNRC6A-containing apical granules represented a potentially novel class of granules distinct from P-bodies in size, subcellular distribution and absence of enzymes needed for mRNA degradation.

### microRNAs are crucial components of the TNRC6A apical granules during multiciliogenesis

We examined the possibility that microRNAs previously reported in MCCs could be locally enriched where the TNRC6A apical granules were abundant. Indeed, ISH for miR-449a, miR-34a, miR-34b, representative MCC miRNAs^23–26^, showed strong signals apically, adjacent to the base of multicilia (**Fig. 3c, Supplementary Fig. 2**). Thus, the data suggested that the TNRC6A apical granules contained AGO2, miRNAs and presumably their targets.

We then asked whether microRNAs were necessary for formation of these apical granules during MCC differentiation. Expression of the miR-34/449 family is required for basal body maturation and apical docking during multiciliogenesis^23^. We examined ALI airway epithelial cultures from miR-34/449 null mice in which all three loci encoding miR-34/449 were deleted^24^. Analysis of ALI day 4 cultures in WT differentiating MCCs showed the typical abundant TNRC6A apical granules revealed by human index serum IF (18033 single-labeled **Fig. 3d, left**) and the larger randomly distributed double-labeled P-bodies (18033-DCP1A-labeled granules). By contrast, in miR-34/449 null cultures, apical TNRC6A granules were notably decreased, absent or disorganized in contrast to the P-bodies, which were largely preserved (**Fig. 3d, right**). Given the high abundance of miR-34/449 in MCCs (~70% of all miRNAs)^23^, the data strongly suggest that miR-34/449 is a key cellular component required for the formation of apical granules. The failure to form these granules is likely linked to events that lead to the organization of the apical domain of differentiating MCCs to initiate multicilia formation.

### Newly-synthesized proteins are concentrated within or adjacent to TNRC6A-containing apical granules

Our study identified TNRC6A, AGO2 and miRNAs in apical granules lacking enzymes for mRNA decay in differentiating MCCs. We asked whether TNRC6A could be involved in mRNA localization and local translation at the apical domain in these cells, which is a site of active multicilia formation. Puromycin has been reported to label newly-synthesized proteins, since it mimics aminoacyl-tRNA and can enter the ribosome A site to terminate translation by ribosome-catalyzed covalent incorporation into the nascent polypeptide C-terminus^27^. O-propargyl-puromycin (OPP), an alkyne analog of puromycin, has been used to detect localized translation^27–29^.

To visualize newly-synthesized polypeptides in differentiating MCCs, ALI Day 4 cultures were labeled with OPP for 30min before termination and subsequently analyzed by IF-confocal microscopy. OPP staining revealed newly-synthesized proteins strongly localized in apical foci underneath emerging cilia (**Fig. 4a**). The OPP foci were cell type-specific and dynamic, as they were not detected in mature MCCs or in other airway epithelial cell types. Notably, OPP signals were abolished by pretreatment with cycloheximide (**Fig. 4b**), supporting the specificity of this approach. OPP foci were similarly found in differentiating MCCs from human ALI airway epithelial cultures, suggesting potential conservation of this mechanism in mammalians (**Fig. 4c**). These apical OPP foci varied considerably in size even within the same MCC In both human and mouse airways (**Fig. 4d**). Morphometric analysis showed OPP foci ranging from 300-2000nm in diameter and with signal intensity about 2.7 folds higher than in their neighboring area (OPP mean optical density, P<0.0001). The marked variation in size and signal intensity suggested a dynamic and heterogeneous regulation of protein synthesis locally in these OPP foci and in their vicinity.

**Fig. 4.**
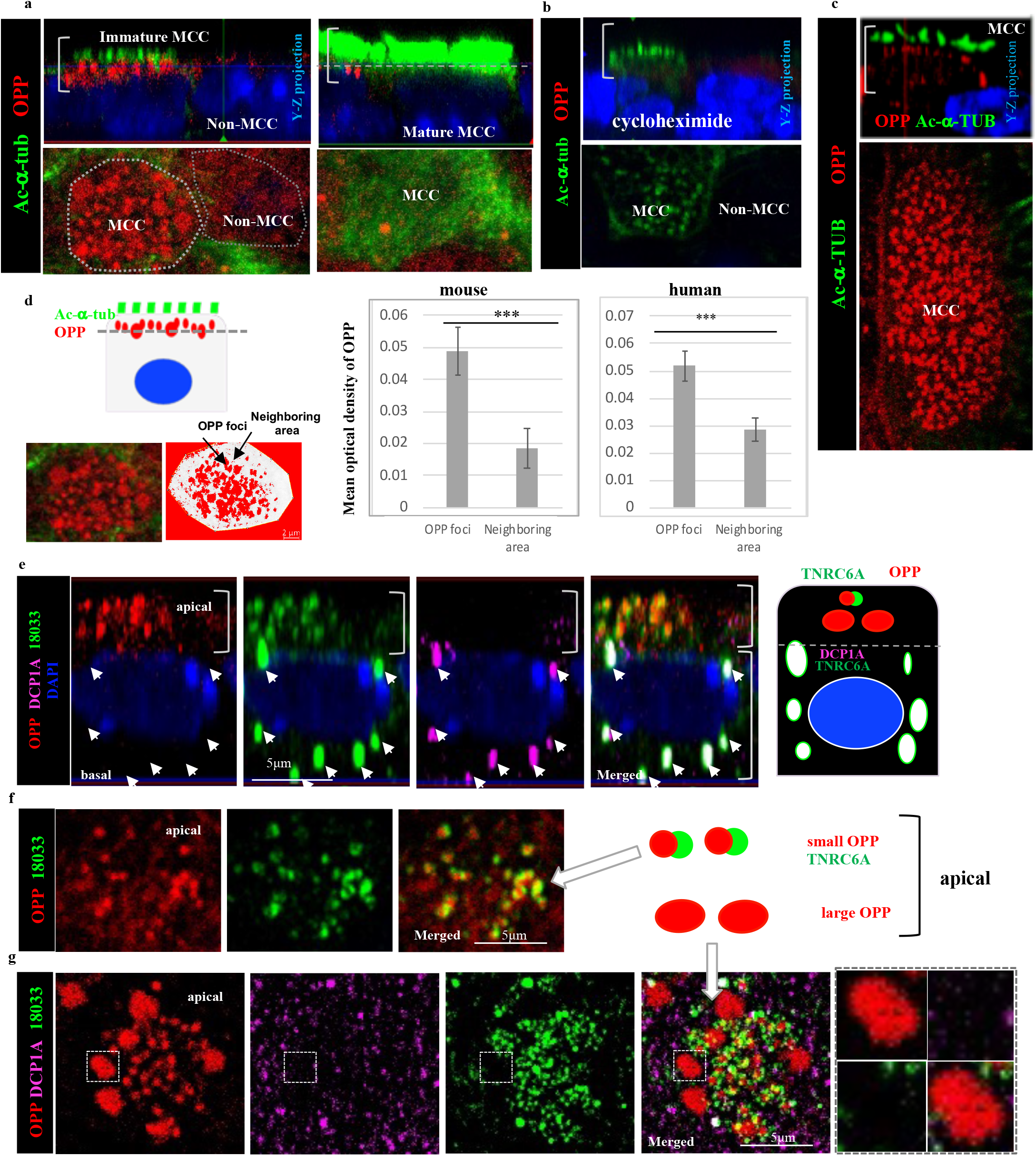
Newly synthesized proteins (OPP) are concentrated within or adjacent to apical TNRC6A granules in ALI day 3 cultures. a) Strong OPP signals in the apical region (brackets) of immature **mouse** MCCs with shorter and less cilia. No OPP signals are present in non-MCCs or in fully differentiated mature MCCs (abundant tall cilia). b) OPP signals are abolished in ALI cultures with cycloheximide, a protein synthesis inhibitor. c) Similar pattern of OPP distribution in apical (brackets) **human** immature MCCs. d) Quantitation of mean optical density of OPP signals in apical granules compared to the neighboring area in the same plane of section (diagram on the left) using Image J software. Bars and lines are mean (± sd) of measurements in 30 immature MCCs from three independent mouse (c) or human (d) ALI cultures ***:P<0.0001. e-g). OPP co-staining with DCP1A and human index serum (18033) showing OPP in the apical MCC region (e, bracket) enriched in TNRC6A granules. No OPP in DCP1A positive P-bodies of the same MCC (e). Heterogeneity of OPP in small and large apical foci (diagrams in e,f). TNR6A granules overlap totally or partially with the smaller apical OPP foci (f) but not with the apical large OPP foci (g). DCP1A signals do not overlap with apical TNRC6A or OPP. f and g are representative Z stack images of the apical region of an immature MCC. Boxed area in g is enlarged on the right panel

Co-staining of OPP with the human index serum (18033) and DCP1A showed OPP signals overlapping with the apical TNRC6A granules but not with the DCP1A positive P-bodies within the same MCC **(Fig. 4e**). Interestingly, within the MCC apical domain, TNRC6A granules overlapped extensively and at various degrees with the smaller OPP foci but not with the larger OPP foci, suggesting TNRC6A-independent events in these larger foci **(Fig. 4f-g and diagrams**).

### Newly-synthesized proteins and translation machinery are found in apical granules during centriole amplification

The proximity of the OPP foci to apically-located ciliary structures suggested an association with subcellular components implicated in multicilia formation ^30,31^. Indeed, co-immunostaining of OPP with markers of these components in ALI day 3-4 cultures revealed co-localization of the smaller OPP foci with DEUP1, a marker of deuterosomes (~400-800nm in diameter)^30,31^, and abutting centrioles (300-400nm in diameter), as identified by Centrin3. The larger OPP foci colocalized with PCM1, which marked fibrous granules (~800-2000nm in diameter)^30,31^ (**Fig. 5a-d**). The spatial relationship of TNRC6A granules with centriole biogenesis components was further confirmed by co-immunostaining of human index serum with Centrin3, γ-tubulin, PCM1, Deup1 (**Supplementary Fig. 3a-d**) and CCP110 (**Fig. 5e)**. The CCP110 signals were found surrounding OPP/DEUP1 double positive foci, suggesting the co-localization of TNRC6A positive granules with procentrioles surrounding deuterosome (**Fig. 5d-e**). Altogether the data strongly suggested a connection between TNRC6A granules and the process of centriole amplification.

**Fig. 5.**
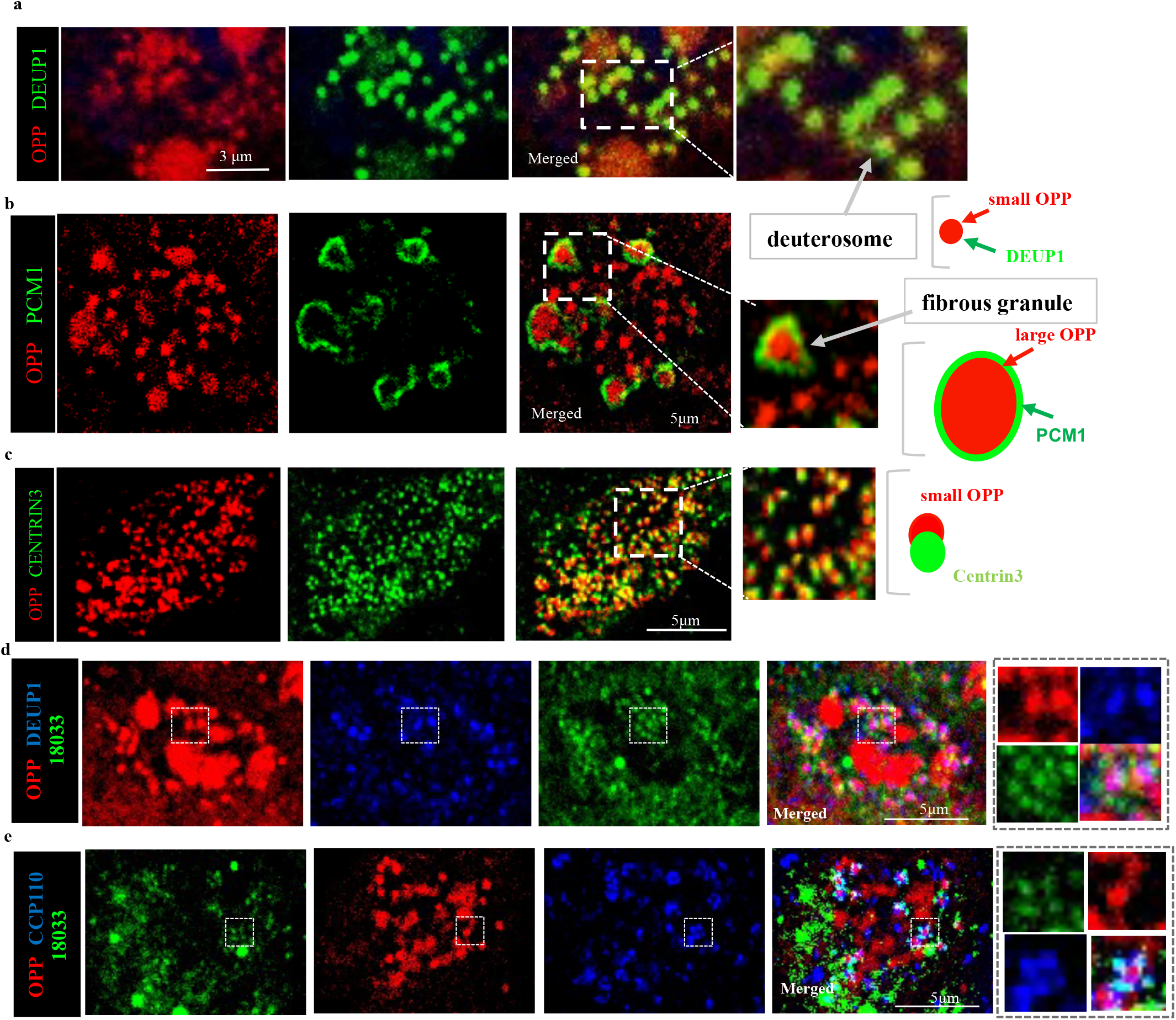
Sites of newly-synthesized proteins are found in apical granules adjacent to centriolar components and fibrous granules in differentiating MCCs. a-c) Strong OPP signals in deuterosomes (DEUP1 positive) and fibrous granules (PCM1 positive), and adjacent or overlapping centrioles (Centrin3 positive). Diagrams depicting the relationship of these components with the small and large OPP foci. d-e). TNRC6A positive granules surrounding OPP foci co-localize with procentrioles marked by CCP110. Deuterosome-associated TNRC6A granules are OPP positive. A plane of Z-stack image of apical region of single MCC in all images. Boxed areas are enlarged on the right panels.

We then asked whether components of the translation machinery were present in the apical OPP-enriched foci. Co-immunostaining of OPP with the translation initiation factors eIF3B or phosphorylated RPS6 (p-RPS6) showed strong signals in the smaller OPP apical granules of differentiating MCCs (**Fig. 6a,b)**. p-RPS6 and eIF3B expression did not overlap with the larger OPP foci but was found in their immediate neighborhood surrounding these foci (**Fig. 6c,d**). Notably, single-molecule fluorescence in situ hybridization (smFISH) for 18S rRNA and 28s rRNA showed strong signals surrounding the large OPP foci (**Fig. 7a,b)**. Using a similar approach we found ribosomal protein 3 (*Rpl3*) mRNA expressed in the small OPP foci characteristic of the apical TNRC6A-containing granules and expressed to a lesser extent in the large OPP foci. MLS mRNA (RNA transcribed from the mitochondrial L-strand DNA), although known to be abundantly expressed in differentiating MCCs, was neither co-localized nor clearly associated with any OPP foci of the ALI day 4 cultures (**Fig. 7c,d)**.

**Figure 6.**
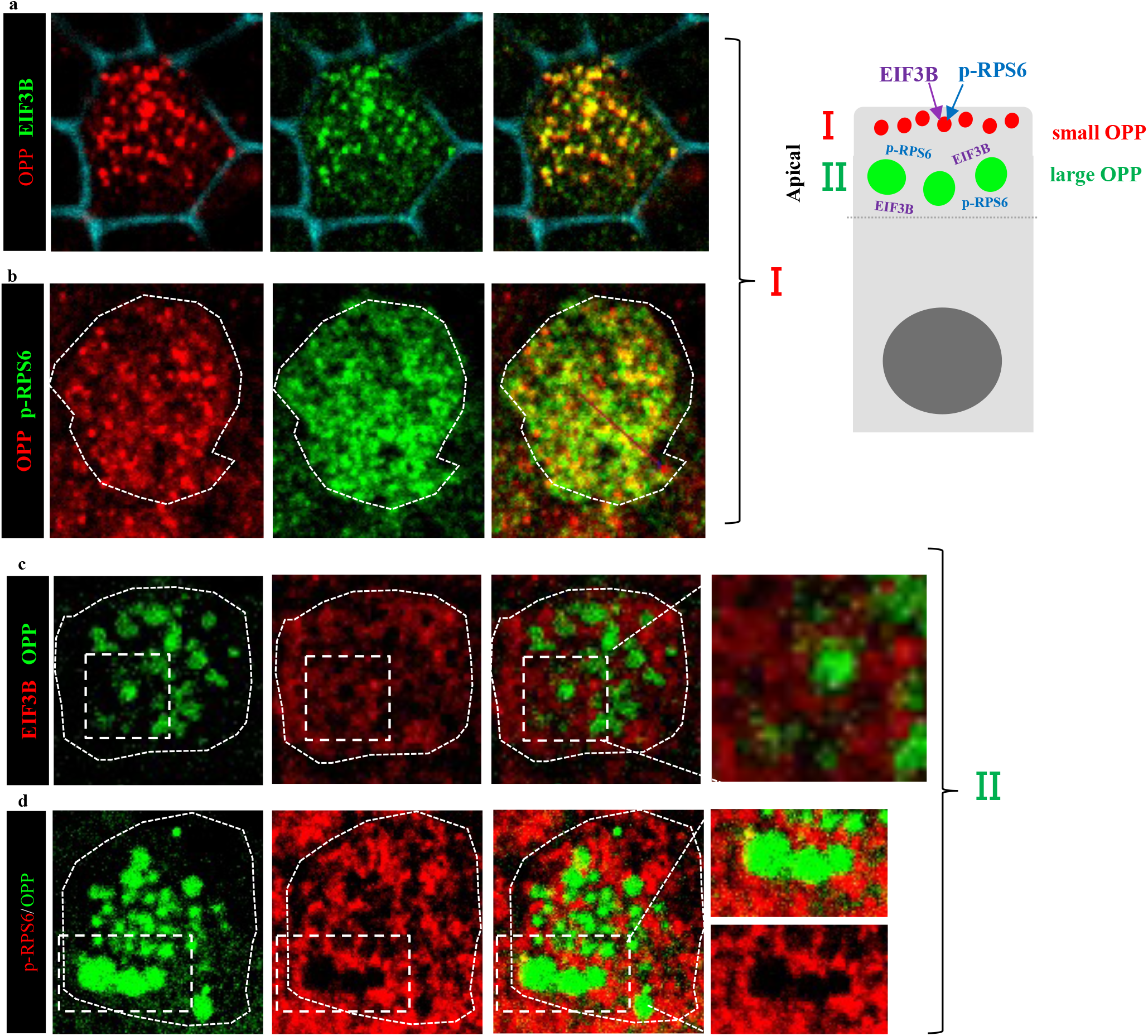
Translation initiation factors are co-expressed or adjacent to OPP foci in differentiating MCCs. All panels. confocal images of a single MCC in ALI day 4 culture. Diagram (right) summarizing the findings. a, b) Translation initiation factors (EIF3B and p-RPS6) overlap with smaller OPP apical foci (**region I**, shown to be TNR6A enriched in Fig.5). Phalloidin-stained cell membrane (blue) depicts apical MCC region. c, d) EIF3B and p-RPS6 proteins are found surrounding the large OPP apical foci (**region II**). High magnification panels on the right enlarged from the boxed areas.

**Fig. 7.**
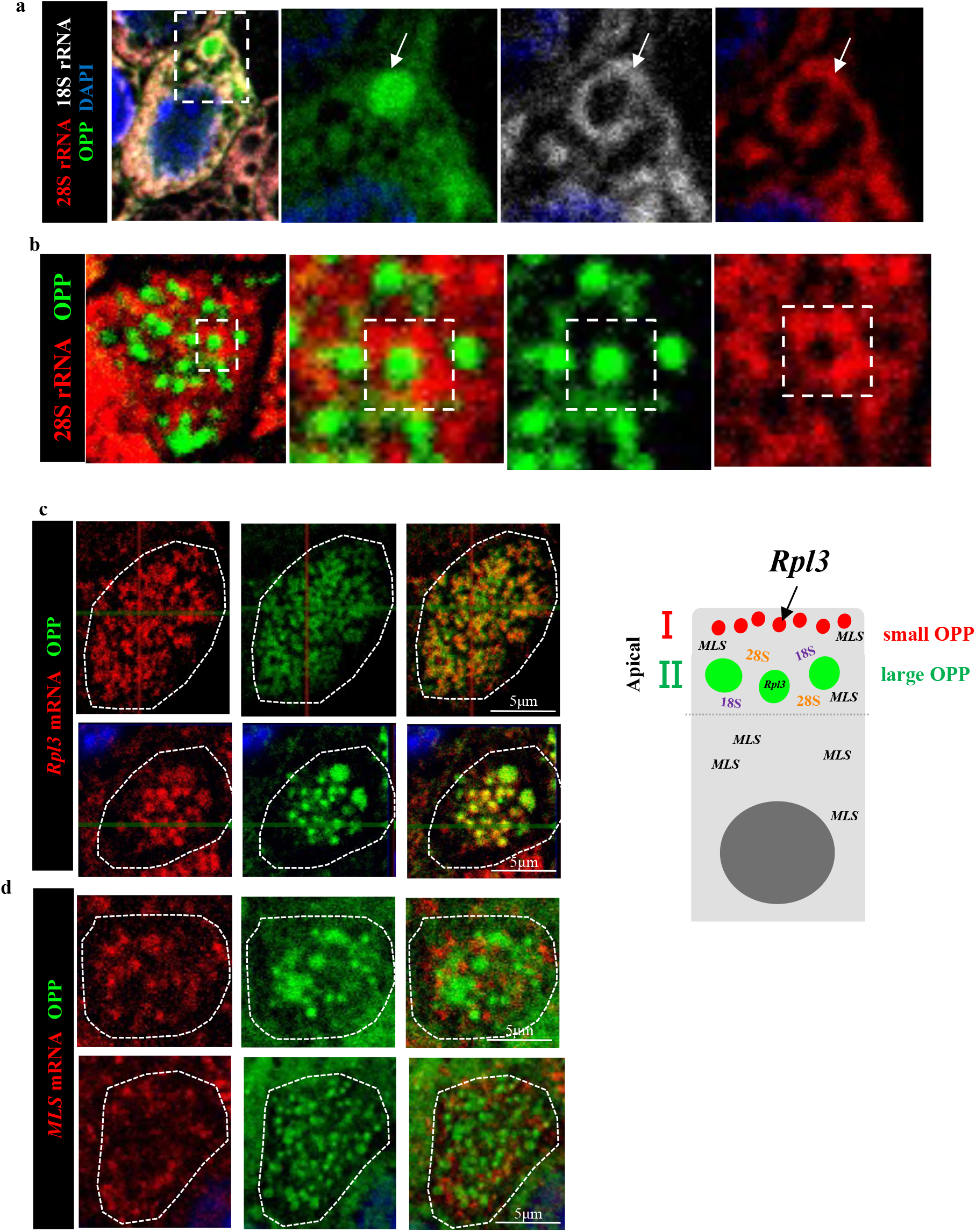
rRNAs and mRNAs for ribosomal proteins are associated with OPP positive foci in differentiating MCCs. RNAs detected by smFISH. A plane of Z-stack image of the apical region of a single MCC shown in each panel.. a-b) 18s and 28s rRNAs (arrows) strongly expressed surrounding OPP foci. Right panels enlarged from boxed areas. c) *Rpl3* mRNA signals strongly present in OPP apical small foci (top panel, shown to be Tnr6a-containing granules in Fig. 4) and expressed to a lesser extent in the large OPP foci (bottom panel) d) RNA transcribed from the mitochondrial L-strand DNA does not overlap with any OPP foci. Diagram summarizing findings (right): apical region I: small OPP foci; apical region II: large OPP foci.

Altogether the findings above suggested that TNRC6A apical granules are active sites of local translation and likely involved **i**n key events in differentiating MCCs.

### Disruption of *Tnrc6a* expression leads to reduced CCP110, abnormal centriologenesis and defective multicilia formation

Next we investigated whether *Tnrc6a* expression was required for the formation of the apical granules in differentiating MCCs. For this, we examined ALI airway epithelial cultures from a *Tnrc6a* null mice also carrying an EGFP reporter to identify cells committed to MCC fate (*Tnrc6a*^*null*^; *FOXJ1-EGFP*, see Methods). IF staining of ALI day 4 with human index serum showed that disruption of *Tnrc6a* specifically abolished the apically-localized granules, but not the cytoplasmic P-bodies in FOXJ1-EGFP-labeled MCCs **(Fig. 8a)**. We asked how this could affect production of CCP110, a key centriolar protein involved in ciliogenesis and known target of miR-34/449^24^. Thus, we co-cultured airway progenitors from wild type and *Tnrc6a*^*null*^; *FOXJ1-EGFP* mutants and differentiated these cells in ALI cultures. This allowed to compare potential differences in the intensity of signals between *Tnrc6a*-deficient and WT cells side by side, using the latter as internal controls. IF analysis of CCP110 showed strong signals in WT cells but expression was drastically reduced in the *Tnrc6a*^*null*^; *FOXJ1-EGFP* MCCs **(Fig. 8b)**. The presence of FOXJ1-EGFP in *Tnrc6a* mutant cells, suggested no significant defect in MCC fate determination (**Fig. 8a-b**). This was further supported by analysis of E18.5 lungs showing that the proportion of FOXJ1 positive cells in airways from WT and *Tnrc6a*^*null*^ mutants was essentially the same (**Supplementary Fig. 4)**.

**Fig. 8.**
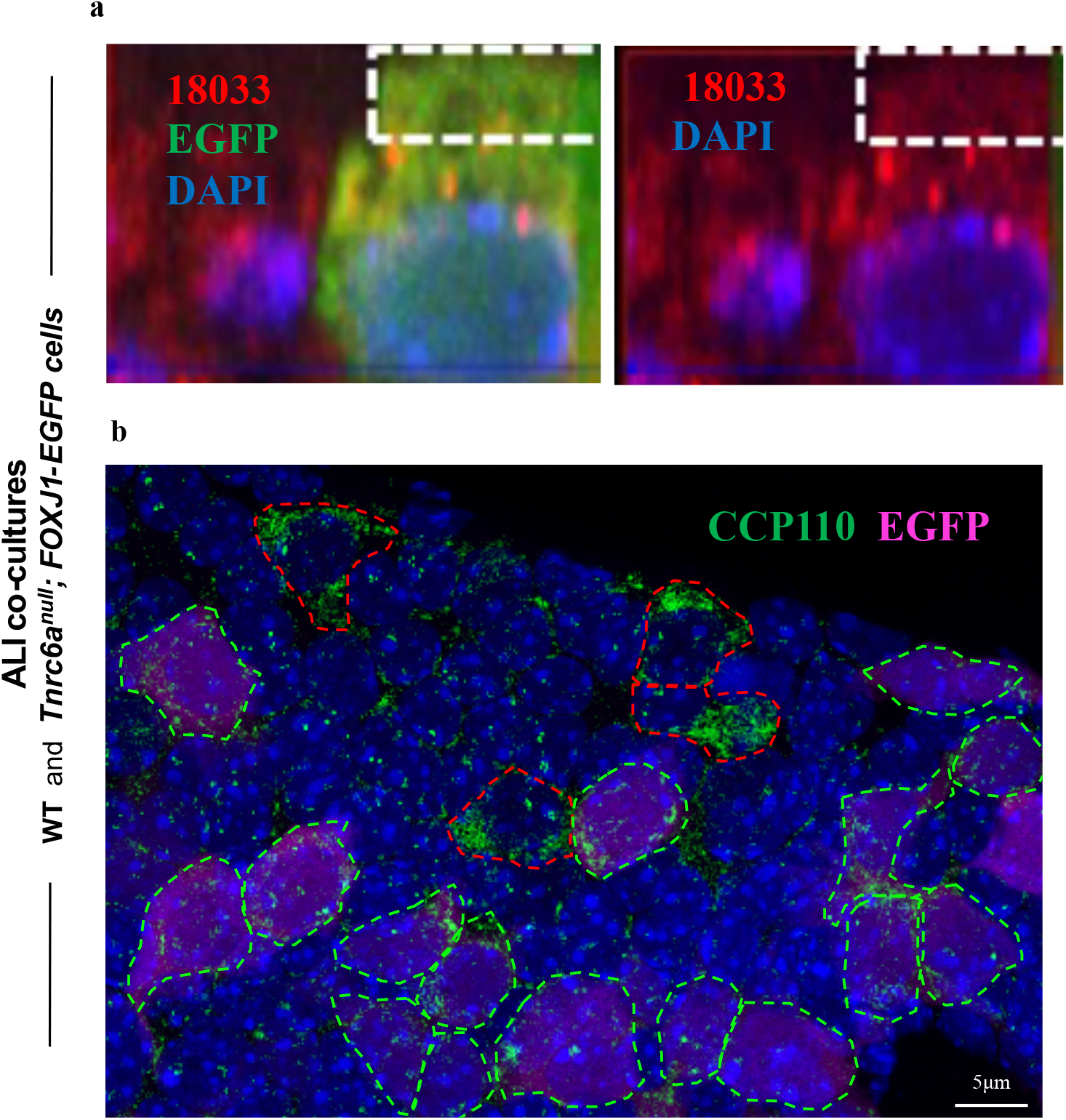
*Tnrc6a* loss of function results in loss of apical granules and reduced CCP110 expression. ALI day 4 co-cultures of WT and *Tnrc6a*^*null*^; *FOXJ1-EGFP* tracheal epithelial cells. a) *Tnrc6a* null mutant MCC marked by FOXJ1-EGFP (green). Human index serum (red) stains both apical granules and P-bodies. However in this mutant MCC apical granules are absent due to loss of Tnr6a (boxed area). b) Strong IF CCP110 signals detected in WT differentiating MCCs (outlined by red dashed line), but nearly absent in *Tnrc6a* null mutant cells (marked by FOXJ1-EGFP, outlined by green dashed line).

IF staining of ALI co-cultures of *Tnrc6a*^*null*^; *FOXJ1-EGFP* and WT epithelial cells revealed marked differences in levels and distribution of proteins involved in centriole amplification. For example, in *Tnrc6a* null cells signals for CEP135 and PCM1 were clustered in a corner of the differentiating MCC, characteristic of an early stage of centriole amplification (**Fig. 9a-b**)^32,33^. Moreover, in mutant MCCs Centrin3 staining was drastically reduced (**Fig. 9c**). These changes were compatible with a significant defect of *Tnrc6a* mutant cells in generating the large number of normal centrioles/basal bodies required for multiciliogenesis. Consistent with a defect in centriologenesis, few cilia formed in differentiating MCCs of *Tnrc6a* mutants when co-cultured with WT cells **(Supplementary Fig. 5a)**. The defect in multicilia formation was further confirmed by examining the trachea of *Tnrc6a* null mutants at E18.5 **(Supplementary Fig. 5b-d)**. Efficient disruption of *Tnrc6a* was demonstrated by IF staining with the monoclonal antibody (4B6), which showed TNRC6A positive granules in tracheal MCCs of wild type but not of mutants (**Supplementary Fig. 5b)**. Co-staining with Ac-α-tub and TNRC6A demonstrated a significantly reduced multicilia staining in *Tnrc6a* null tracheas. Scanning electron microscopy of tracheae from *mutants* showed the majority of MCCs (83%) with shorter and sparser cilia as compared to WT (**Supplementary Fig. 5c, d)**. The cilia truncation was consistent with an immature MCC phenotype. We found no difference in FOXJ1 staining between WT and *Tnrc6a* null lungs (**Supplementary Fig.4**), suggesting no defect in MCC fate determination. Transcriptome analysis of WT and *Tnrc6a* null lungs did not detect significant changes in expression of genes associated with proliferation or differentiation of other cell types of the lung. Moreover, no gross abnormalities were observed in these embryos or in the overall morphology of their lungs and no compensatory increased expression of other *Tnrc6* family member (*Tnrc6b* and *Tnrc6c*) was observed in Tnrc6A null lungs (**Supplementary Fig. 6**)^18^.

**Fig. 9.**
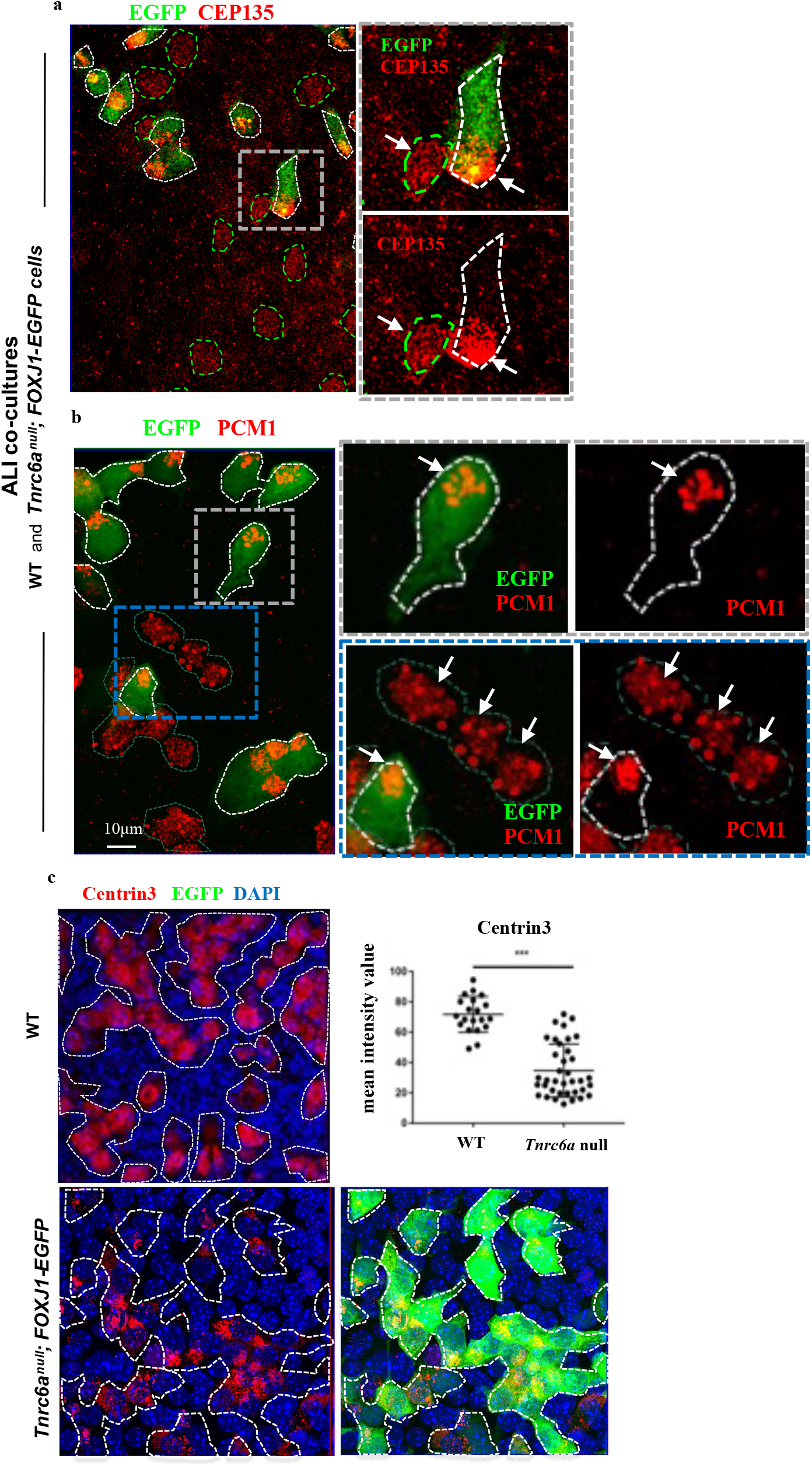
Defective centriole amplification in *Tnrc6a-*deficient cells. a, b) ALI day 4 co-cultures of WT and *Tnrc6a*^*null*^; *FOXJ1-EGFP* tracheal epithelial cells. IF for CEP135 (a) or PCM1 (b): Asymmetric distribution of signals (arrows) clustered in the corner of *Tnrc6a* null defective MCCs (marked by FOXJ1-EGFP, white dashed) contrasting with the homogeneous signal distribution in WT (green dashed) MCCs. Boxed areas enlarged in the right panels. c) IF of Centrin3 protein in ALI day 4 culture of WT (top panel) and *Tnrc6a null -* FOXJ1-EGFP tracheal epithelial cells (bottom panels). Dashed lines depict differentiating MCCs in WT and mutants (visualized in green by FOXJ1-EGFP). Marked decrease in Centrin3 signals in mutant MCCs confirmed by quantitation of signal intensity (Image J software). Graph: dot plot (mean, ± sd of 30 MCCs from three independent experiments). Student’s Test;. ***: P<0.0001.

Together, these data strongly support that idea that loss of *Tnrc6a* leads to a defect selectively in MCCs, the same cell type where we found *Tnrc6a* highly expressed in WT lungs.

### Disruption of Tnrc6a expression leads to reduction, rather than stabilization of miRNA targets in mutant lungs

TNRC6A has been traditionally described as a component of the RNA-induced silencing complex in microRNA (miRNA)-mediated gene suppression, repressing translation or promoting mRNA degradation. This is consistent with previous findings of defective miRNA function with increased miRNA target expression in yolk sac endoderm of Tnrc6a null mice^18^.We examined the subcellular localization of TNRC6A in E9.5 yolk sacs and found endodermal signals colocalized with AGO2, EDC4 and other mRNA degradation enzymes in randomly-distributed cytoplasmic granules suggestive of P-bodies^17^ (**Supplementary Fig. 7**). Intriguingly, our analysis of TNRC6A expression in the lung showed evidence of its localization in differentiating MCCs in apical granules distinct from P-bodies in size, subcellular distribution, absence of enzymes needed for mRNA degradation and enrichment in translation machinery components. Moreover we had evidence of TNRC6A apical granules as sites of active translation crucial for large-scale centriole amplification in multiciliogenesis. This suggested that TNRC6A could have a distinct role in the lung, compared to the yolk sac, where it promotes miRNA target degradation. To investigate this possibility we examined the behavior of miRNA in the lung and yolk sac from control and Tnr6a null mice.

We performed gene set enrichment analysis (GSEA) using a dataset containing the most evolutionary conserved targets of 83 conserved miRNAs as determined by Targetscan (http://www.targetscan.org). GSEA of these genes in yolk sac (GSE30244) showed targets of most miRNAs (78 out of 83) significantly enriched in the list of genes upregulated in *Tnrc6a* null yolk sacs (**Supplementary Fig. 8**). This was consistent with a role for *Tnrc6a* in the destabilization of miRNA targets^18,34^. Strikingly, similar analysis using the *Tnrc6a* null lung dataset (GSE89327) showed that targets of most miRNAs (79 out of 83) were enriched among the genes significantly reduced in *Tnrc6a* mutant lungs, but not among the genes upregulated (**Supplementary Fig. 9)**. This was consistent with the idea that, in the lung, *Tnrc6a* stabilizes miRNA targets instead of promoting degradation.

Indeed, analysis of miR-34/449 predicted targets in both yolk sac and lung datasets showed that, among 94 transcripts with conserved binding sites significantly altered in *Tnrc6a* null lungs, the majority (80%) was downregulated. By contrast 75% of the conserved miR-34/449 targets altered in yolk sac mutants were upregulated (72 total). Among the top 30 most downregulated genes in mutant lungs, several have been implicated in multicilia formation and 30% of these genes are predicted targets of miR-34/449, key in multiciliogenesis^23,24^ (**Supplementary table 1, Supplementary Fig. 10**). For example, Spef2 and Rsph1 were strongly expressed in MCCs of the developing lung and their signals were reduced in *Tnrc6a*^null^ mutants. qRT-PCR further confirmed the reduced expression of additional miR-34/449 targets, such as *Rnf213, Tmem159* and *Calcrl*, besides *Rsph1, Spef2* (**Supplementary Fig. 10**). As demonstrated in other systems, genetic deletion of *Tnrc6a* disrupted miRNA function without affecting miRNA expression or stability^18^. Analysis of miR-449 (from the miR34-449 cluster) and miR-146 showed no difference in expression between WT and *Tnrc6a* null mutants (**Supplementary Fig. 11)**.

Altogether, our data strongly suggested that, in the lung, *Tnrc6a* was not involved in degradation of miRNA targets but rather appear to be required for mRNA stabilization, and likely promoting protein synthesis.

## Discussion

Here we identified TNRC6A selectively expressed in MCCs of lungs *in vivo*, concentrated in a novel class of apical granules containing AGO2, miRNAs and their targets, adjacent to immature centrioles. Although present in adult airways, these granules are mostly abundant in differentiating immature MCCs. Remarkably, components required for mRNA degradation are missing in these granules, while present in P-bodies within the same MCCs. Instead, they are enriched in components of the mRNA translation machinery and show evidence of concentrated newly-synthesized proteins. We found that, in the lung, loss of *Tnrc6a* leads to broad reduction instead of accumulation of miRNA targets, suggesting a role in target mRNA stabilization, rather than in degradation. Loss of *Tnrc6a* resulted in a drastic reduction of expression of centriolar proteins, known miRNA targets, abnormal centriologenesis and, ultimately, defective MCCs.

Cytoplasmic RNP granules in mammalian cells, including P-body, stress granule, neuronal transport granule and germ granule are generally linked to translation repression. mRNAs are not translated in stress granules where translation initiation factors are concentrated. Indeed, we found no evidence of newly-synthesized polypeptides in P-bodies in differentiating MCCs. Instead, OPP and translational machinery components were present in the apical TNRC6A-containing granules, suggesting that these are sites of active translation. Our findings, together with reports of active translation associated with FUS granules ^35,36^, point to a more prevalent role of cytoplasmic RNP granules in active local translation than previously suspected.

Recruitment of TNRC6A to AGO2/miRNA complex triggers typically a cascade of events for translation suppression and target mRNA degradation^18,37,38^. However, in addition to recruiting deadenylation complexes to trigger mRNA degradation, TNRC6A can also interact with poly (A) binding proteins (PABP)^39,40^ to promote local translation. AGO2 mainly functions as a miRNA binding protein to trigger miRNA-mediated mRNA degradation^41,42^. Interestingly, AGO2 was originally purified and cloned as a translation initiation factor and named EIF2C2^43^. It is known that AGO2 can interact with EIF1A in the biogenesis of miRNAs^44^. miRNAs are well known as a negative regulators of gene expression. However, miRNAs have been also shown to stimulate mitochondrial translation to activate translation of miRNA targets in quiescent cells and to enhance translation initiation mediated by HCV-like IRESes^45–47^.

Recent studies have identified TNRC6A or AGO2 adjacent to the basal body of primary cilia or centrosomes^48,49^, suggesting that the spatial relationship between TNRC6A/AGO2 positive granules and centrioles is conserved in both primary cilia and multicilia (as we report here). We found newly-synthesized proteins (OPP) also concentrated in granules at the base of primary cilia (**Fig. 10**). However, primary cilia remained intact in cultured *Tnrc6a* null cells in contrast to our findings in multicilia. Our data are consistent with a mechanism in which miRNA pathway components (here TNRC6A, AGO2) and miRNAs (such as miR-34/449) in apical granules of differentiating MCCs recruit proteins (PABP, EIFs) and target mRNAs for stabilization and local translation of centriole biogenesis proteins crucial for multiciliogenesis. Proximity-dependent biotinylation (BioID) assays using centriolar and cilia proteins as baits have shown TNRC6A and AGO family members in close proximity to CEP135 and CEP152 ^50^. Here we had no evidence of centriolar proteins associated with enzymes involved in mRNA degradation.

**Fig. 10.**
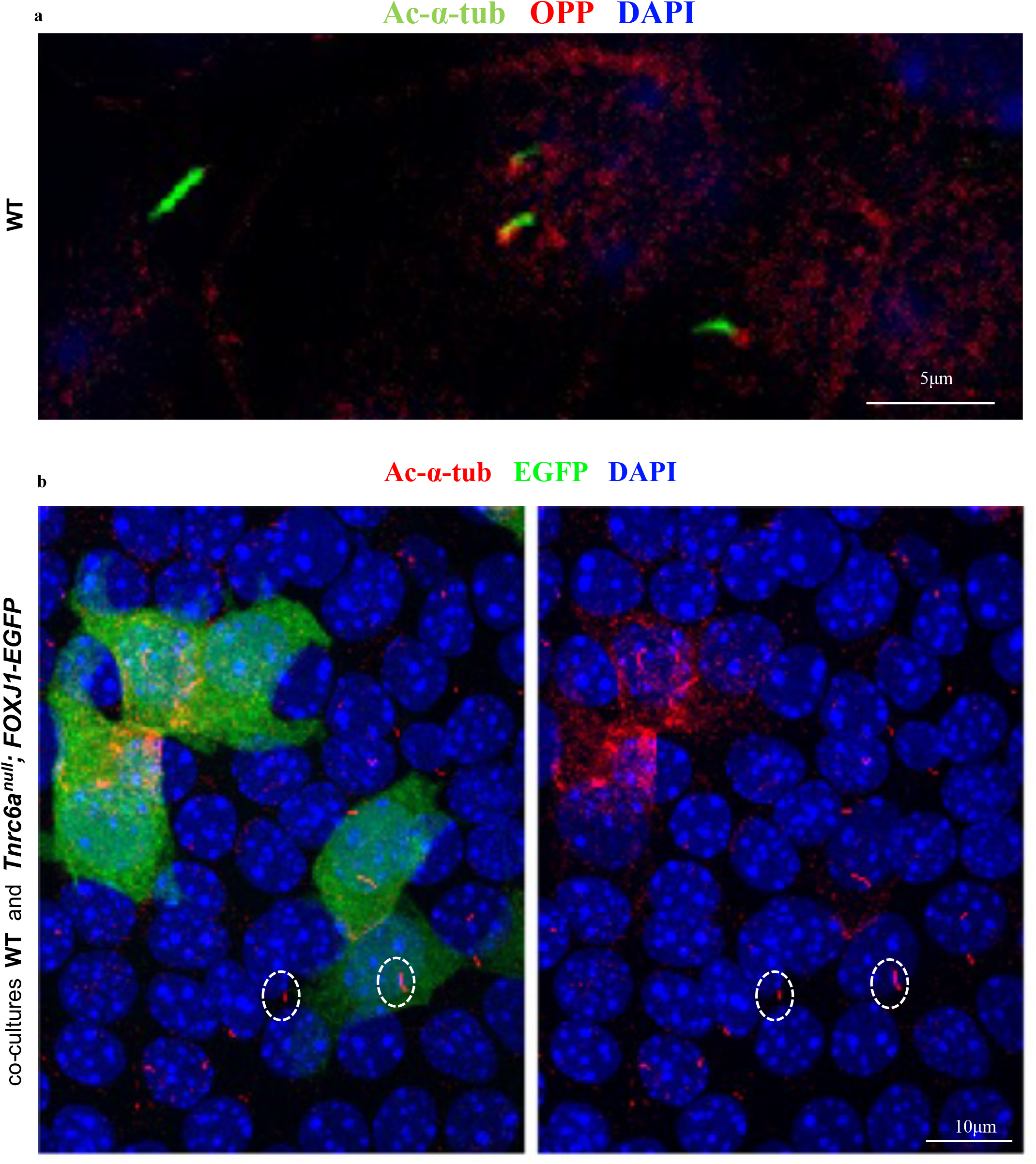
Disruption of *Tnrc6a* does not prevent primary cilia from forming. IF of ALI airway epithelial cultures at day3 a) WT ALI airway epithelial cultures: newly-synthesized proteins (red, label ed by OPP) are concentrated in foci at the base of primary cilia (green). b) Co-culture of WT and Tnrc6a null; FOXJ1-EGFP cells at ALI day3 : *Tnrc6a* null cells marked by FOXJ1-EGFP (green) showing no significant defect in primary cilia marked by Ac-α-tub staining (circled).

In summary, our findings suggest that components of the miRNA pathways can be organized into subcellular hubs for target mRNA localization, stabilization and local translation crucial for differentiation of airway MCCs. These observations further provide strong evidence of the critical role of mRNA localization for efficient delivery and concentration of subsets of proteins to specific intracellular sites for prompt organelle biogenesis during cellular differentiation.

## Acknowledgments

We thank Jun Qian for technical assistance; Dr. Andreas Moor and Dr. Shalev Itzkovitz from the Department of Molecular Cell Biology at Weizmann Institute of Science, Israel, for sharing with us probes for smFISH. We also thank the Lu and Cardoso lab members for thoughtful discussions and input. This study is supported by grants from NHLBI (R01 HL131422 to J.L and W.V.C and R35-HL135834-01 to W.V.C).

## Materials and Methods

### Mouse breeding, genotyping and monitoring

*Tnrc6a*^gt/+^ reporter mice carrying a gene trap insertion of the *β-galactosidase* gene in the *Tnrc6a* locus and Tnrc6a^gt/gt^ mutants were generated on a mixed genetic background as described^18^. Tnrc6a mutants were backcrossed to C57BL/6 for ten generations to overcome embryonic lethality. Mice are housed in a non-barrier animal facility at Columbia University Medical Center. The genotyping is performed as reported^18^. miR-34/449 null mice were bred and genotyped as described^23^. Foxj1/EGFP/Tnrc6A mice were generated by breeding Tnrc6A^gt^/+ mice to Foxj1-EGFP mice (Jax, B6; C3-Tg (Foxj1-EGFP)85Leo/J)^51^. All animal studies were carried out with the approval of the Institutional Animal Care and Use Committee (IACUC), Columbia University Medical Campus.

### Cloning and packaging of lentiviral vectors

pHAGE-EGFP-Tnrc6a vector was cloned by replacing EGFP fragment of pHAGE-CMV-eGFP-W1 vector (generous gift from Dr. Darrel Kotton, Boston University Medical School) with EGFP-Tnrc6a DNA fragment from pT7-EGFP-Tnrc6A (Addgene #25035) plasmid. To do this, we first cut pHAGE-CMV-EGFP-W1 with NotI (NEB # R0189S), followed by the treatment with DNA Polymerase I, Klenow Fragment (NEB # M0210S) to blunt the end. Then EGFP fragment was released by cutting with BamHI (NEB # R0136S), followed by separation and purification of vector fragment for ligation. To prepare EGFP-Tnrc6a fragment, P7-EGFP-Tnrc6a was cut with AgelI (NEB # R0552S), followed by the treatment with DNA Polymerase I, Klenow Fragment (NEB # M0210S) to blunt the end. Then cut again with BamHI to release and purify EGFP-Tnrc6A fragment. DNA sequencing was performed following ligation and transformation to identify and confirm the right clone. Similarly, EGFP-Ago2 fragment was released from EGFP-Ago2 plasmid (EGFP-C1 backbone, addgene, # 21981) by AgeI, blunted and followed by BamH1 digestion. The same fragment of lentiviral vector used in pHAGE-EGFP-Tnrc6A cloning was used for the ligation to generate pHAGE-EGFP-Ago2. Packaging of all lentiviral vectors was performed by co-transfection of 293T cells (80% confluent, 15 cm plate) with 2ml of Trans-IT/DNA/media mix containing 5 plasmids encoding the lentiviral backbone (24ug) as well as tat (1.2ug), rev (1.2ug), gag/pol (1.2ug) and vsv-g (2.4ug) viral genes together with 90μl of Trans-IT (MIR #6600). Viral particles in the resulting supernatants were collected 48hrs after transfection and concentrated by ultracentrifugation (90 minutes at 48960g on Beckman SW28 rotor). Titers used in all experiments range from 0.5-1.5 × 10^9^TU/ml. For the constructs above, lentiviral infection of basal cells isolated from mouse trachea was performed at the time of seeding and the culture were terminated at Day 3 of the differentiation, when the centriologenesis and ciliogenesis are highly active.

### Histological analyses

Tissues were dissected and fixed overnight in in 4% paraformaldehyde (Ted Pella, #18505) overnight at 4°C, then processed either for frozen section or paraffin section by following standard procedures. All the blocks were sectioned at 10 μm either for immunostaining or for in situ hybridization.

### β-galactosidase staining

This was performed according to protocol described previously^18^. Briefly, adult lungs of Tnrc6A^gt/+^ mouse were perfused intratracheally with 0.25% glutaraldehyde (Electron Microscopy Science, #16220) at a constant pressure of 30cm H_2_O for 30min, then perfused with 1XPBS for 2 hours, followed with staining buffer (5μm C6N6FeK3, 5μm C6N6FeK4, 2mM MgCl2, 1mg/ml X-gal in 1XPBS) for 30min. Then the trachea was tied and the whole lungs were incubated in the staining buffer in a 50ml tube overnight at 37°C. After washing in 1XPBS three times (2hours for each wash), lungs were fixed again using 4% PFA (Ted Pella, #18505) for 2 hours and then processed for embedding following standard procedures. Sections of LacZ-stained lungs were either subjected to H&E staining or co-stained with antibodies specific for MCCs.

### In situ hybridization (ISH)

ISH detection of miR-34a and miR-449a and miR-92b or *Tnrc6a* mRNA expression was performed on frozen sections of mouse lungs using 5’ digoxigenin-labeled LNA probes (Exiqon) or digoxigenin-labeled antisense probe of *Tnrc6a*. Briefly, slides were washed with DEPC-treated 1X PBS 3 times for 3 minutes each, followed by the treatment with 100mM triethanolamine buffer (Sigma, Cat #90279) plus 0.25% of acetic anhydride (Sigma, #320102) for 10 minutes, then permeabilized by incubating in PBST (1X PBS plus 0.1% Triton X-100 (Sigma, #9002-93-1) in DEPC-treated water for 30 minutes, and washed 3 times for 5 minutes each at RT in 1X PBS. After 2 hours pre-hybridization using hybridization buffer without labeled probe (50% formamide (American Bioalnalytical, # 95-12-7), 10mM Tris-HCl pH8.0, 600mM NaCl, 1X Denhardt’s solution (Sigma, #02532), 200μg/mL tRNA, 1mM EDTA, 0.25% SDS (Ambion, #AM9823), 10% dextran sulfate (American Bioalnalytical, # ab427), hybridizations were carried out at 55°C overnight in the hybridization buffer containing 25nM of DIG-labeled miR-34a (#38487-01, ACAACCAGCTAAGACACTGCCA) or miR-449a (Exiqon, #38587-15, ACCAGCTAACAATACACTGCCA) or miR-92b (Exiqon, #610819-340, CTGCACCGCGTCCCGTC). Then, slides were sequentially washed with SSC (2X, 1X, 0.2X) at 50°C for 30 min. followed by 0.2X SSC (Ambion, #AM9763) for 5 minutes at RT, and finally 1X PBS for 5 minutes at RT. Slides were then incubated in blocking solution (TTBS, 0.05M Tris, pH 7.5, 0.15M NaCl, 0.1% Tween-20, plus 5% sheep serum) and incubated with anti-digoxigenin-AP, Fab fragments (1:500, Roche, Cat. #11093274910) overnight at 4°C. After washing in TTBS 3 times for 10 minutes each, signals were developed using BM purple (Roche, # 11442074001). Hybridization for Tnrc6A using digoxigenin-labeled antisense probe was performed at 70°C by following the standard procedure. ISH of Spef2 and Rsph1 was performed by RNAscope according to manufacturer’s protocol.

### Immunofluorescence (IF) and immunohistochemistry (IHC)

For IHC, paraffin sections were deparaffinized and rehydrated by sequentially incubating sections in three washes of xylene (5 min each), two washes of 100% ethanol (10 min each), two washes of 95% ethanol (10 min each), followed by two washes in dH_2_O (5 min each). Endogenous peroxidase was quenched by incubation with 0.3% H_2_O_2_/0.3% horse serum in 1X PBS for 5 minutes at RT. Then, slides were washed in TNT buffer (0.1M Tris pH 7.5, 0.15M NaCl, 0.05% Tween-20) 3 times (5min each), followed by blocking with TNB buffer (0.1M Tris pH 7.5, 0.15M NaCl, 0.5% blocking reagent) for one hour at RT. Incubation with Ac-α-tub antibody (Sigma, #T7451, 1:10,000) diluted in TNB buffer overnight at 4°C. Slides were then washed with TNT buffer 3 times, incubated with Horseradish peroxidase (HRP) conjugated secondary antibodies (Life Technologies) for 2 hours at RT, signals were developed by using the DAB staining kit (Vector Laboratories, Cat.# SK-4100). When TNRC6A monoclonal antibody (Santa Cruz, 4B6, # sc-56314) was used for immunohistochemistry, TSA Plus Fluorescence Kit (Perkin Elmer, # NEL741B001KT) was used for signal amplification. After incubating with HRP-conjugated secondary antibody, signals of slides or ALi filters were developed by TSA Plus Working Solution following manufacture’s protocol. For IF staining, cryosections or transwell filter of ALI cultures were washed in PBS, and then blocked by 5% donkey serum in PBST (1XPBS, 0.3% Triton X-100) for one hour at RT, followed by incubation with primary antibodies overnight at 4°C (TNRC6A, human index serum, 1:3000; FOXJ1, eBioscience, # 14-9965-82, 1:100; Ac-α-tub, Sigma, # T7451, 1:10,000, or Abcam # ab125356; DCP1A, Abcam, #ab183709, 1:50; DDX6, Cell Signaling Technology, #8988S, 1:20; EDC3, Santa Cruz, # sc-55081, 1:20; EDC4/GE-1, Cell Signaling Technology, #2548S, 1:50; Xrn1, Bethyl Laboratories, #A300-443A, 1:20; γ-tubulin, Abcam, #ab84355, 1:50; EIF3B, Santa Cruz, # sc-16377, 1:50). DEUP1, Dr Xueliang Zhu’s lab, 1:300; PCM1, Cell Signaling, #5213, 1:50. Alexa Fluor 488 or 568-conjugated secondary antibodies (Donkey anti-Mouse IgG, or Donkey anti-Rabbit IgG or Donkey anti-Goat IgG; Life Technologies, 1:250 dilution) and Alexa Fluor 594 phalloidin (Life Technologies, #A22287) were used for the visualization of signals. Zeiss 710 confocal microscope was used for the image acquisition and analysis.

### Differentiation of MCCs in Air-Liquid Interface (ALI) culture of tracheal progenitors

ALI culture of mouse tracheal epithelial cells (mTEC) were performed using basal cells isolated from adult mouse tracheas as described previously^21^. Briefly, tracheas from adult (8- to 12-weeks old) mice were dissected and cleaned in ice-cold Ham’s F-12 medium (Life Technologies, #31765-092) with 1% penicillin/streptomycin (Life Technologies, # 15140-163). To isolate epithelial cells, tracheas were digested in 1.5mg/ml Pronase (Roche, #10165921001) in Ham’s F-12 media overnight at 4°C. The digestion was stopped by fetal bovine serum (FBS, sigma, #12133C, a final concentration of 10%). The epithelial cells including basal cells were dislodged from trachea by gentle inverting the tube, and medium containing epithelial cells was transferred to another tube. This was repeated for three time and medium containing epithelial cells were pooled and Centrifuge at 500Xg, 4°C, for 10 min. After removing the medium, epithelial cells were treated with DNase (200μl per trachea) on ice for 5min, and spin down again at 500Xg, 4°C. The pelleted cells were washed by pre-warmed mTEC/Basic medium containing advanced DMEM/F-12 (Life Technologies, #12634-028), 15mM HEPES (Life Technologies, #15630-080), 1% penicillin/streptomycin, 4mM glutamine (Life Technologies, #35050-061), 3.6mM sodium bicarbonate (Life Technologies, #25080-094), and 0.25ug/mL Amphotericin B (Life Technologies, #15290-018), and centrifuged at 500 × g for 5 min. Then cells were resuspended in mTEC/Plus medium, which contained mTEC/Basic medium with 5% FBS, 25ng/ml Epidermal Growth factor (EGF) (Corning, #354001), 10μg/ml Insulin (Sigma, #I6634), 5 μg/ml Transferrin (Sigma, #T1147), 0.1 μg/ml Cholera Toxin (Sigma, #C8052), 30 μg/ml Bovine Pituitary Extract (BPE)(sigma,#P1476), and freshly added 50nM retinoic acid (RA, Sigma, #R2625), and plated on rat tail type I collagen (Advanced Biomatrix, #5056) coated trans-wells (Corning, #3470) of 24-well plates. 10uM Rock inhibitor Y27632 (Tocris, # 1254) was added to mTEC/Plus only once when the cells were plated. The mTEC/Plus medium was changed every other day for 7 days, until we switched the culture to air-liquid interface culture by removing the media in the apical chamber, and replacing the media in the basal chamber with mTEC/SF medium. The mTEC/SF medium was changed every other day until the culture was terminated at different time point. Cultures were terminated at different time point after the induction of differentiation and harvested for RNA isolation or Immunofluorescence. For IF, cultures filters were washed with room temperature PBS twice, then fixed by 4% PFA for 15min, followed by three time of washes in PBS, and keep them in PBS at 4°C.

### Single-molecule fluorescence in situ hybridization (smFISH)

mTEC epithelial cultures were fixed for 15 min in 4% PFA, followed by three washed in PBS then transfer to 70% ethanol for storage. In situ hybridization and imaging were performed as previously reported^28^. Fluorescence labeled DNA oligo probes are from Itzkovitz’s Laboratory. All probes are labeled with Cy5 except rRNA 18s (TMR).

### O-Propargyl-puromycin (OPP) labeling and imaging

mTEC cultures were treated with either 50μg/ml Cycloheximide (Sigma-Aldrich, C7698) or DMSO control for 15 min, followed by 30 min incubation with 50μM OPP (Jena Bioscience, NU-913-5). Cultures were then washed with room temperature PBS and fixed with 4 % of PFA, followed by three washes in PBS. OPP features an alkyne group that can be detected and imaged by the addition of azide-coupled fluorophores in the highly specific click reaction^27,28^. The click reaction was performed subsequently to the IF staining by making use of the Click-iT Alexa Fluor 488 or Click-iT Alexa Fluor 647 Imaging Kits (Thermo Fisher Scientific, C10337 and C10340). The kits were used according to the manufacturer’s instruction with the exception of an additional 20-fold dilution of the working solution of the Alexa Fluor azide dye in DMSO.

### Immunofluorescence (IF) signal quantitation and Statistical analysis

To quantify the IF staining of newly synthesized proteins (OPP) and other centriolar proteins, Z-stack of confocal images were taken for the entire cells. For OPP staining, several planes covering the entire OPP foci were chosen and the maximal projection images of these planes were acquired. Then the areas encompassing the OPP foci and background were marked using ImageJ and the mean optical intensity was calculated. For centriolar proteins, we first generated the maximal projection image for entire cells, and drew the cell boundaries. Then the mean optical intensities were measured by ImageJ. Data were presented as the mean ± SEM and subjected to 2-tailed, paired Student’s t test. P < 0.05 was considered significant. In all experiments a minimum of 3-4 replicates (ALI cultures or lung sections) were used for analyses.

### Scanning Electron Microscopy (SEM)

Tracheas from E18.5 WT or Tnrc6A^gt/gt^ embryonic lungs were dissected, fixed in 2.5% glutaraldehyde, 0.1 M sodium Cacodylate, 0.2 M Sucrose, 5mM MgCl2 pH 7.4 and dehydrated through a graded series of ethanol dilutions. Samples were processed using liquid carbon dioxide in a Tousimis Samdri 795 Critical Point Drier (Rockville MD), sputter coated with chromium in a Quorum EMS 150T ES (Quorum Technologies Ltd, United Kingdom) and examined in a Zeiss Supra Field Emission Scanning Electron Microscope (Carl Zeiss Microscopy, LLC North America), using an accelerating voltage of 2 KV. For quantitative analysis, lungs (n=2) were examined and about 120 cells were counted for each sample.

## Data availability

Data referenced in this study from transcriptomics of control and Tnr6a null mice in lung (E18.5) and yolk sac (E9.5) are available in the National Center for Biotechnology Information Gene Expression Omnibus (GEO) and are accessible through the GEO Series accession number GSE89327, GSE30244 and GSE89327. The data is available in link https://drive.google.com/drive/folders/15JZ_G_kxdZSpjS9dw1tIHfZEds-dUNwk?usp=sharin. All other relevant data are available from the corresponding author upon request.

## Legends

### SUPPLEMENTARY FIGURES

**Supplementary Fig. 1.**
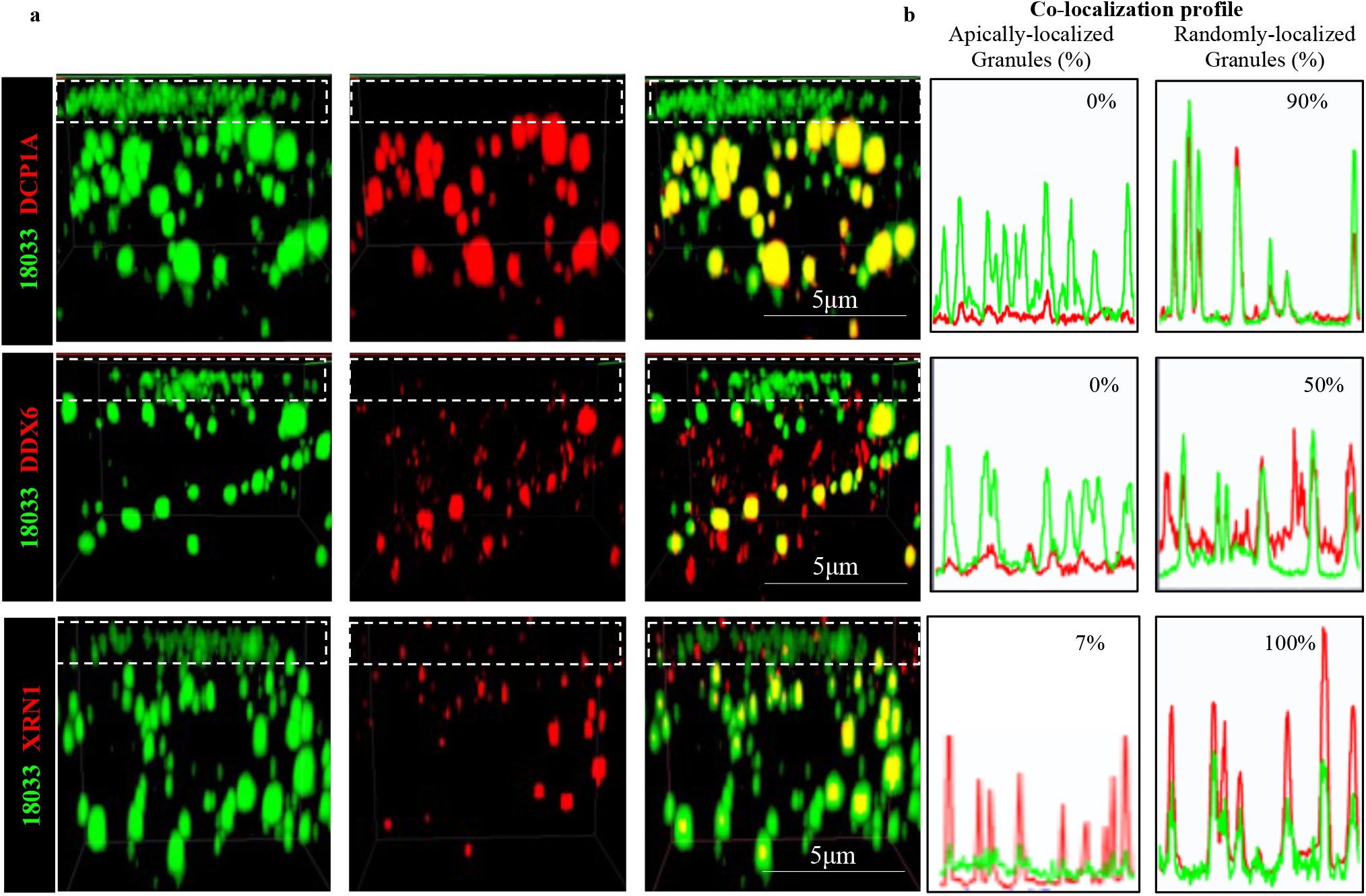
TNR6A-containing apical granules labeled by 18033 lack enzymes for mRNA degradation. a) Each panel represents a 3D image of confocal Z-stack images of a single MCC from an ALI day 3 culture. Apical granules are highlighted by dashed lines. TNRC6A stained with human Index serum (18033). DCP1A, DDX6 and XRN1 are concentrated in randomly-localized granules (P-bodies). However, DCP1A and DDX6 are undetectable in apically granules. XRN1 is present in small number of apically granules at low levels. b) Co-localization profile of apical or randomly-localized granules (P-bodies) within the same MCC indicating the expression of these proteins in the two classes of granules and the percentage of granules with double-labeling signal.

**Supplementary Fig. 2.**
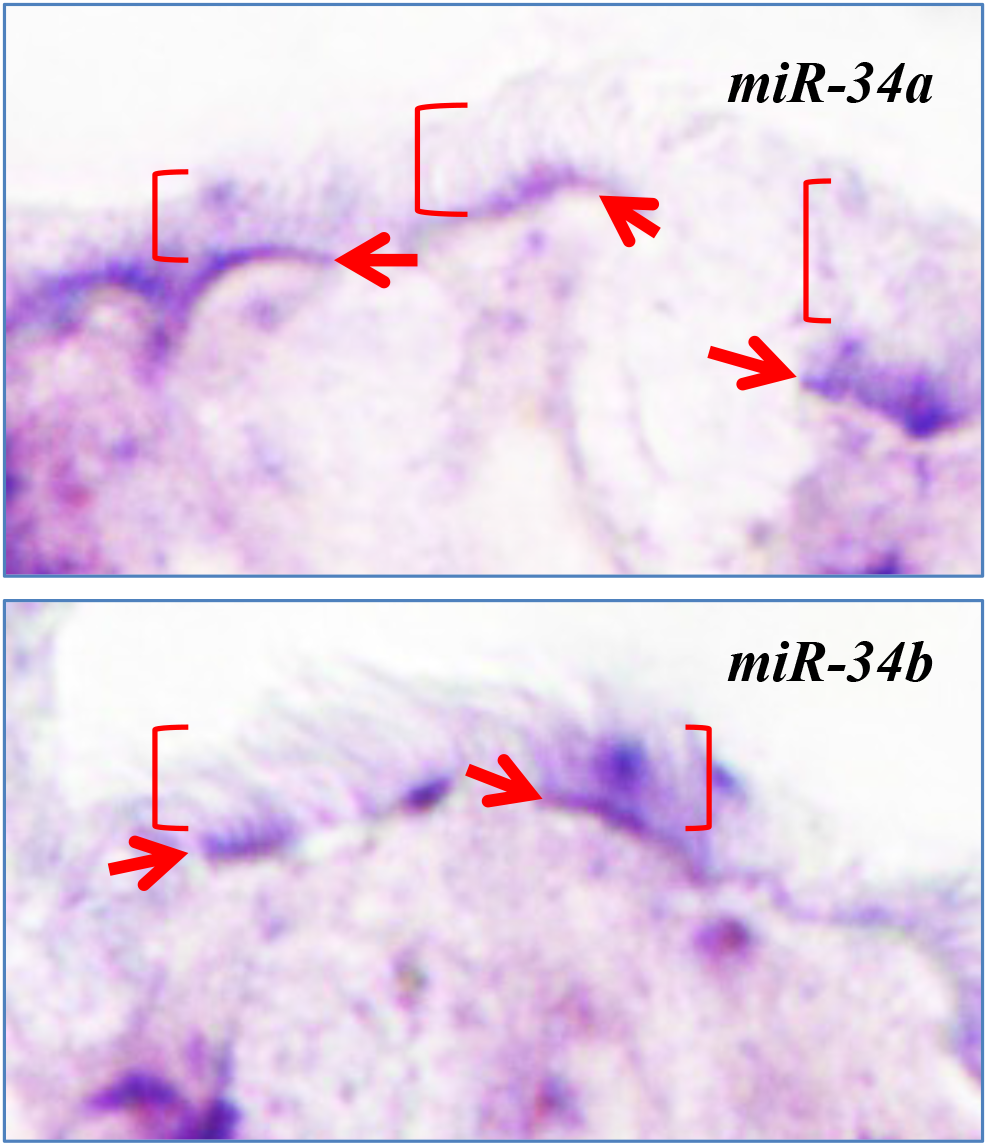
Apically-enriched ISH signal of miRNAs in multiciliated cells of airways in vivo. a-b) In situ hybridization of miR-34a and miR-34b. Apically-localized signals (arrows) underneath multicilia (bracket)

**Supplementary Fig. 3.**
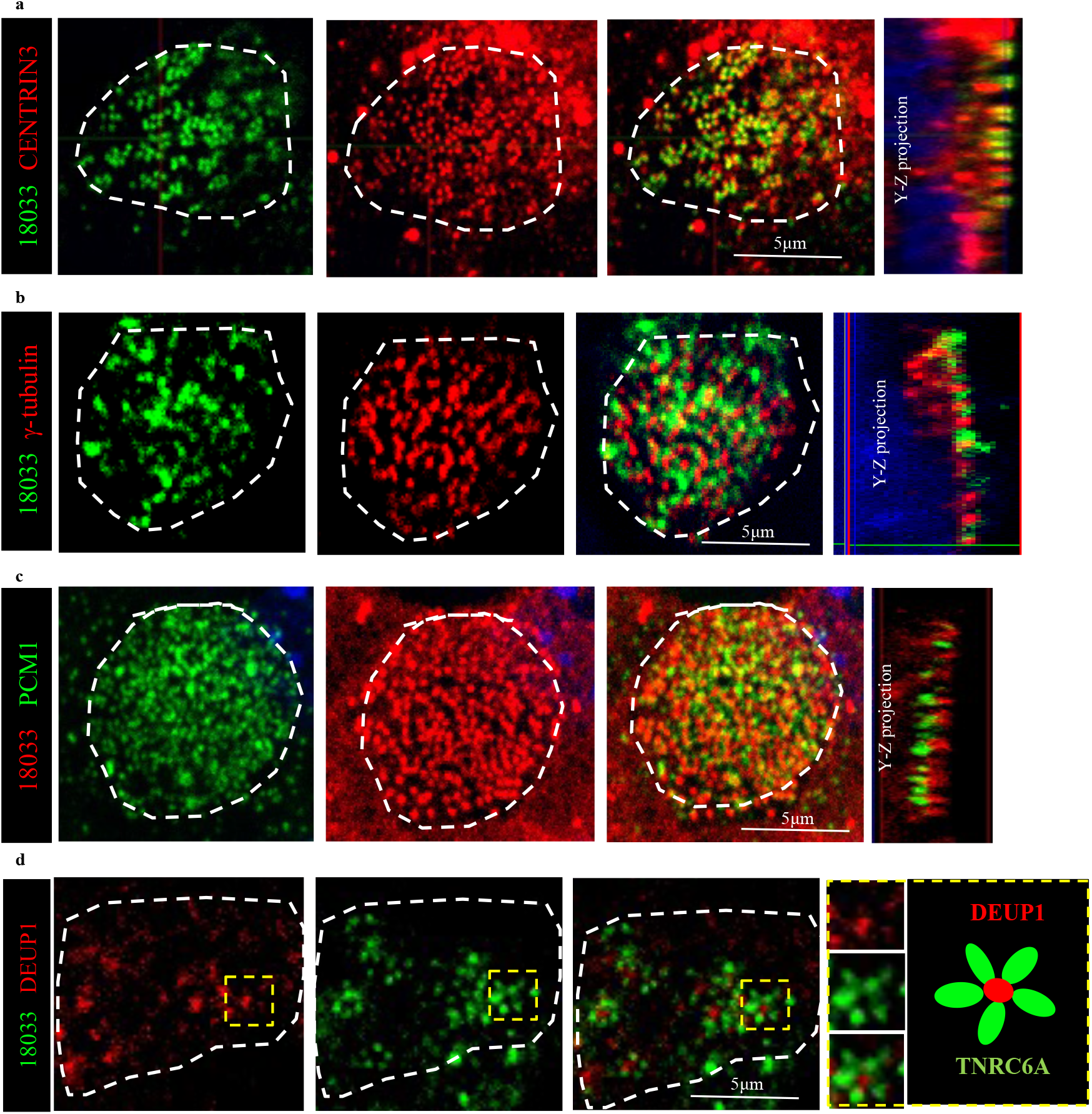
TNRC6A-containing apical granules are adjacent to centrioles in differentiating MCC. Co-staining of TNRC6A (green or red, human index serum, 18033) with Centrin 3 (red, a), γ-tubulin (red, b), PCM1 (green, c) and DEUP1 (red, d). Images in each panel are representative of a plane of Z-stack of the apical region in a single MCC in ALI culture.

**Supplementary Fig. 4.**
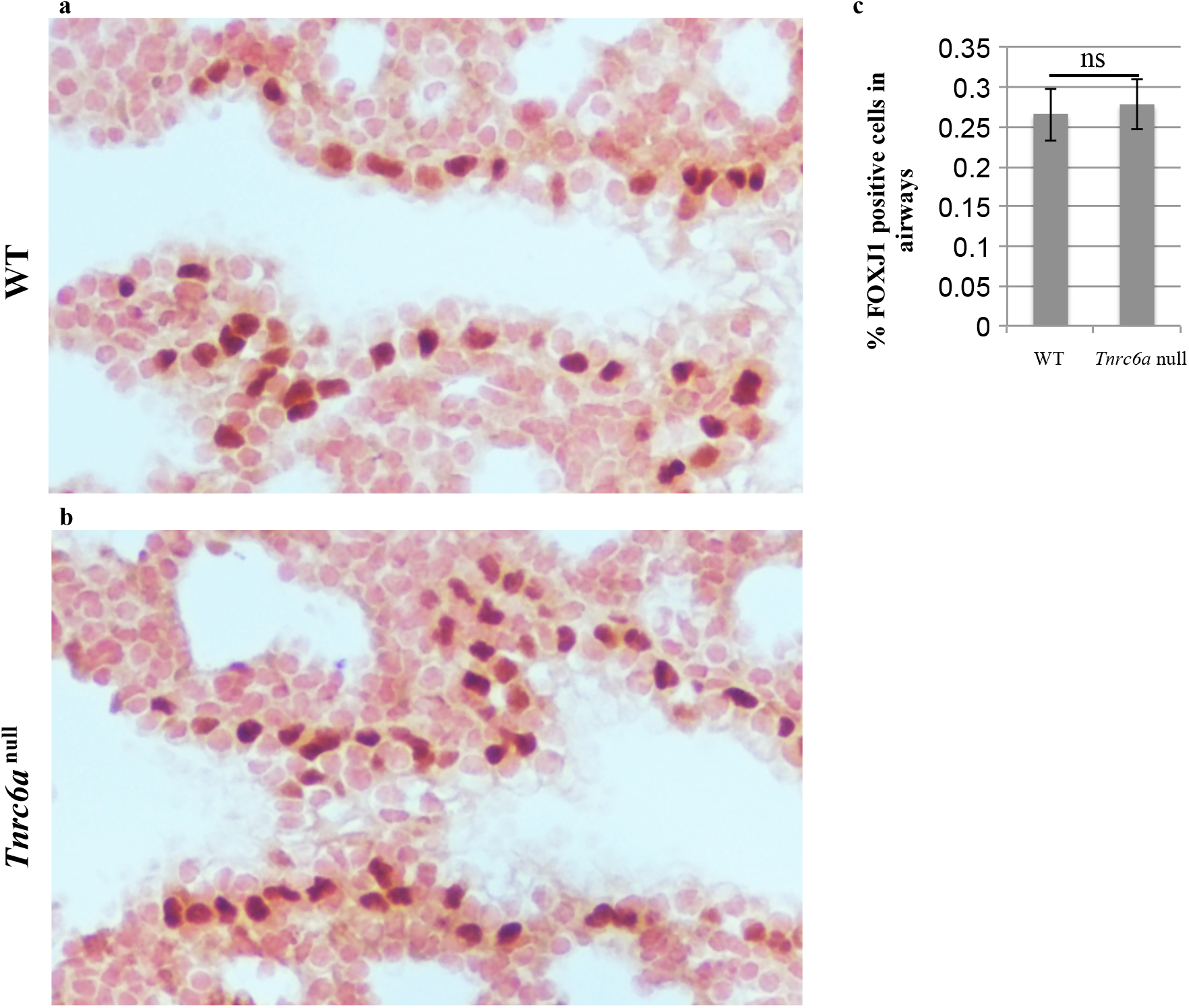
No differences in FOXJ1 expression between wild type and *Tnrc6a null* airways. a-b) FOXJ1 staining in airways of E18.5 wild type and *Tnrc6a* null lungs. c) Percentage of FOXJ1 positive cells in the airways of wild type and *Tnrc6a* null embryos. Bars are mean ± SEM. Student’s T test. ns: non-significant

**Supplementary Fig. 5.**
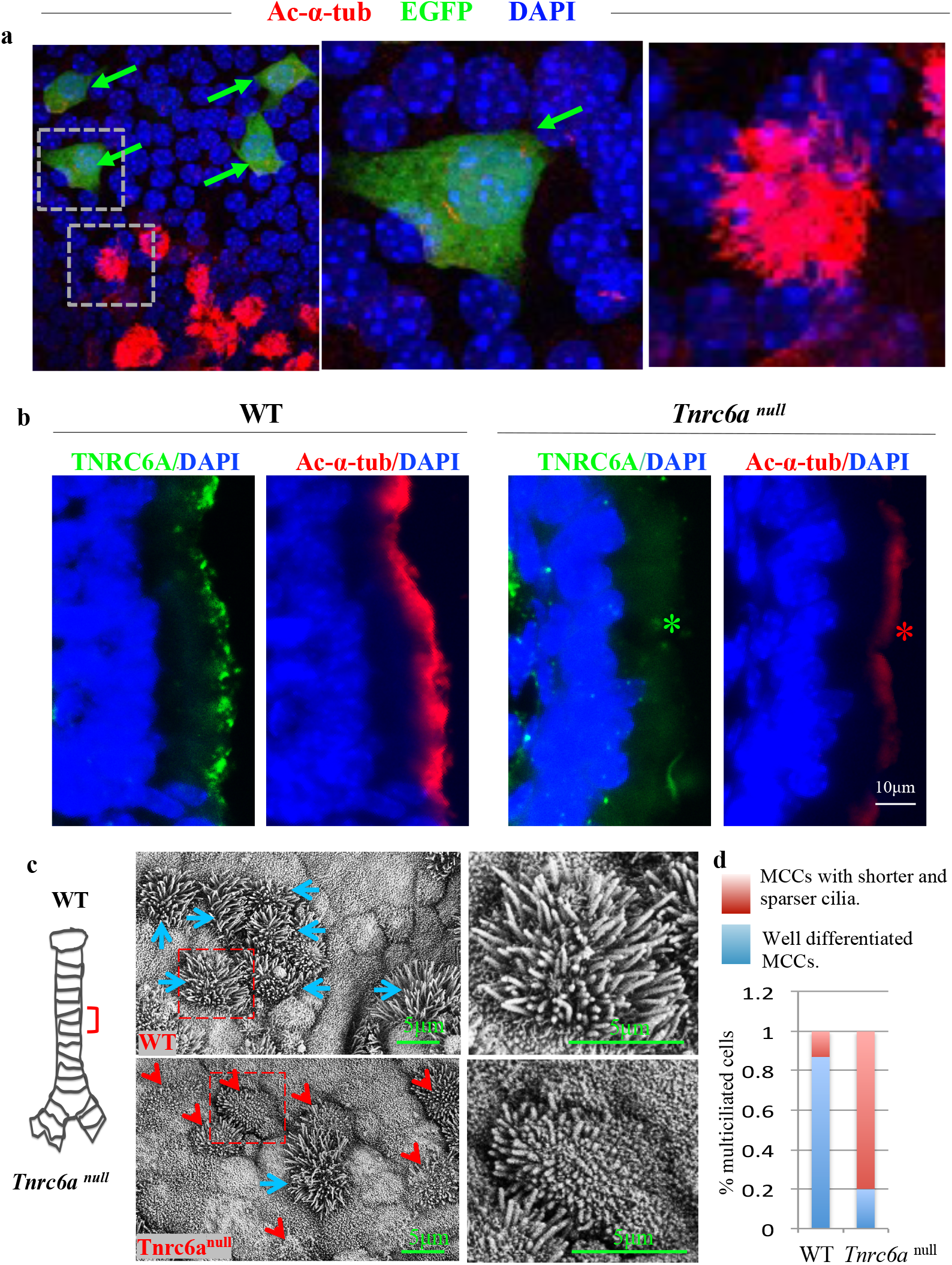
Disruption of *Tnrc6a* results in defective multicilia formation. a) ALI day 4 co-cultures of WT and Tnrc6a null;FOXJ1-EGFP tracheal epithelial cells. Tnrc6a mutant cells (arrows, green) fail to form multicilia (marked by Ac-α-tubulin). Boxed area enlarged on the right panels. b) Co-IF staining of TNRC6A (green, monoclonal antibody 4B6) and Ac-α-tub (red) of wild type or *Tnrc6a*^*null*^ E18.5 airways. Major decrease in Ac-α-tub signals in mutants (*) suggestive of defective multiciliogenesis. c) Scanning electron microscopy of E18.5 trachea of wild type or *Tnrc6a* null mice (right panels are enlarged images of highlighted areas by dashed lines); defective (red arrowhead) and well differentiated (blue arrow) MCCs in WT and mutant trachea. d) Percentage of MCCs with short and sparse cilia in WT and *Tnrc6a* null tracheas (n=2, p<0.01). a) ALI day 4 co-cultures of WT and Tnrc6a null; FOXJ1-EGFP tracheal epithelial cells. Tnrc6a mutant cells (green) fail to form multicilia (marked by Ac-α-tubulin). Boxed area enlarged on the right panels. b) Co-IF staining of TNRC6A (green, monoclonal antibody 4B6) and Ac-α-tub (red) of wild type or *Tnrc6a*^*null*^ E18.5 airways. Major decrease in Ac-α-tub signals in mutants (*) suggestive of defective multiciliogenesis. c) Scanning electron microscopy of E18.5 trachea of wild type or *Tnrc6a* null mice (right panels are enlarged images of highlighted areas by dashed lines); defective (red arrowhead) and well differentiated (blue arrow) MCCs in WT and mutant trachea. d) Percentage of MCCs with short and sparse cilia in WT and *Tnrc6a* null tracheas (n=2, p<0.01).

**Supplementary Fig. 6.**
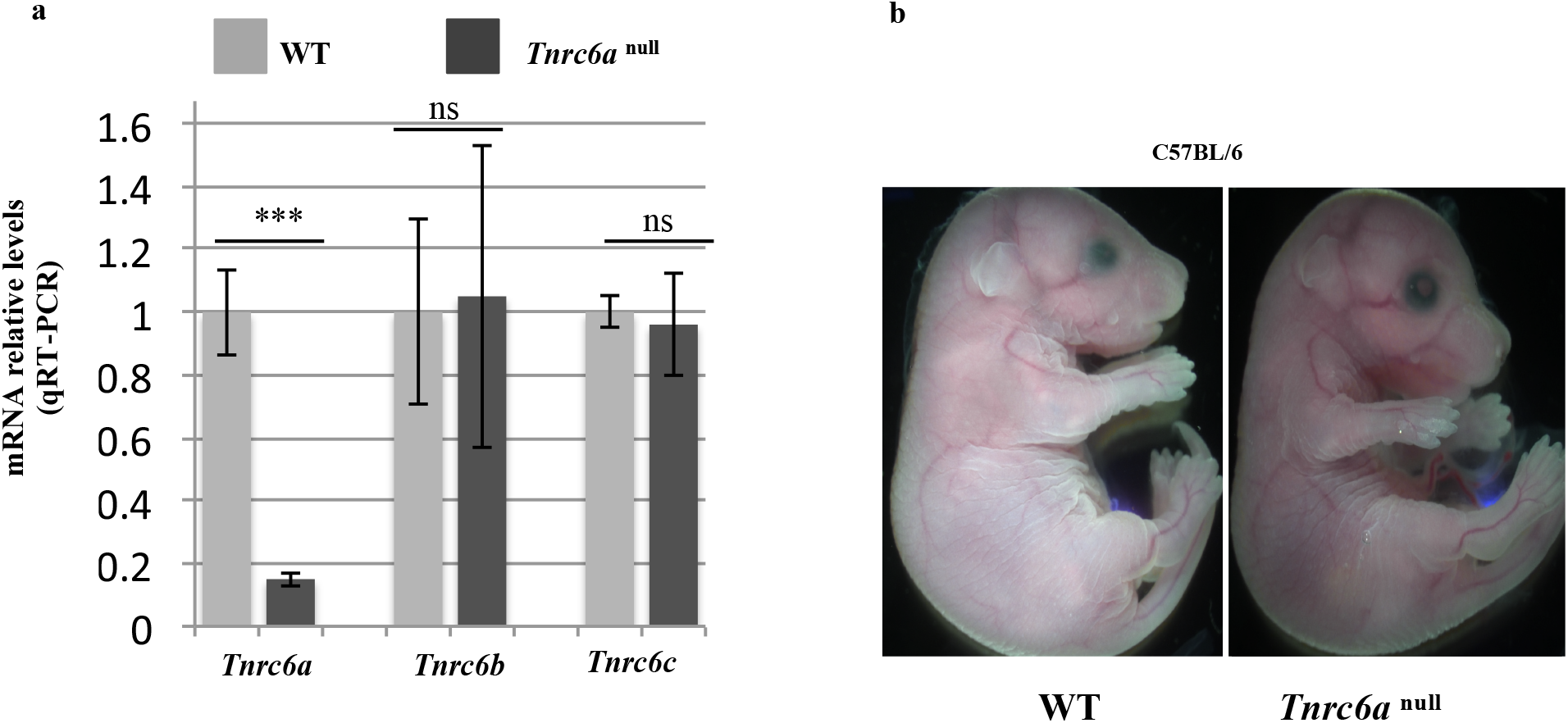
a) Disruption of *Tnrc6a* expression in *Tnrc6a* ^null^ lungs results in no compensation by *Tnrc6b* and *Tnrc6c* expression. qRT-PCR. Graph: Bars are mean ± SD of n=3 lungs per group. b) No gross abnormalities were observed in *Tnrc6a* ^null^ embryos at E18.5 compared to WT littermates.

**Supplementary Fig. 7.**
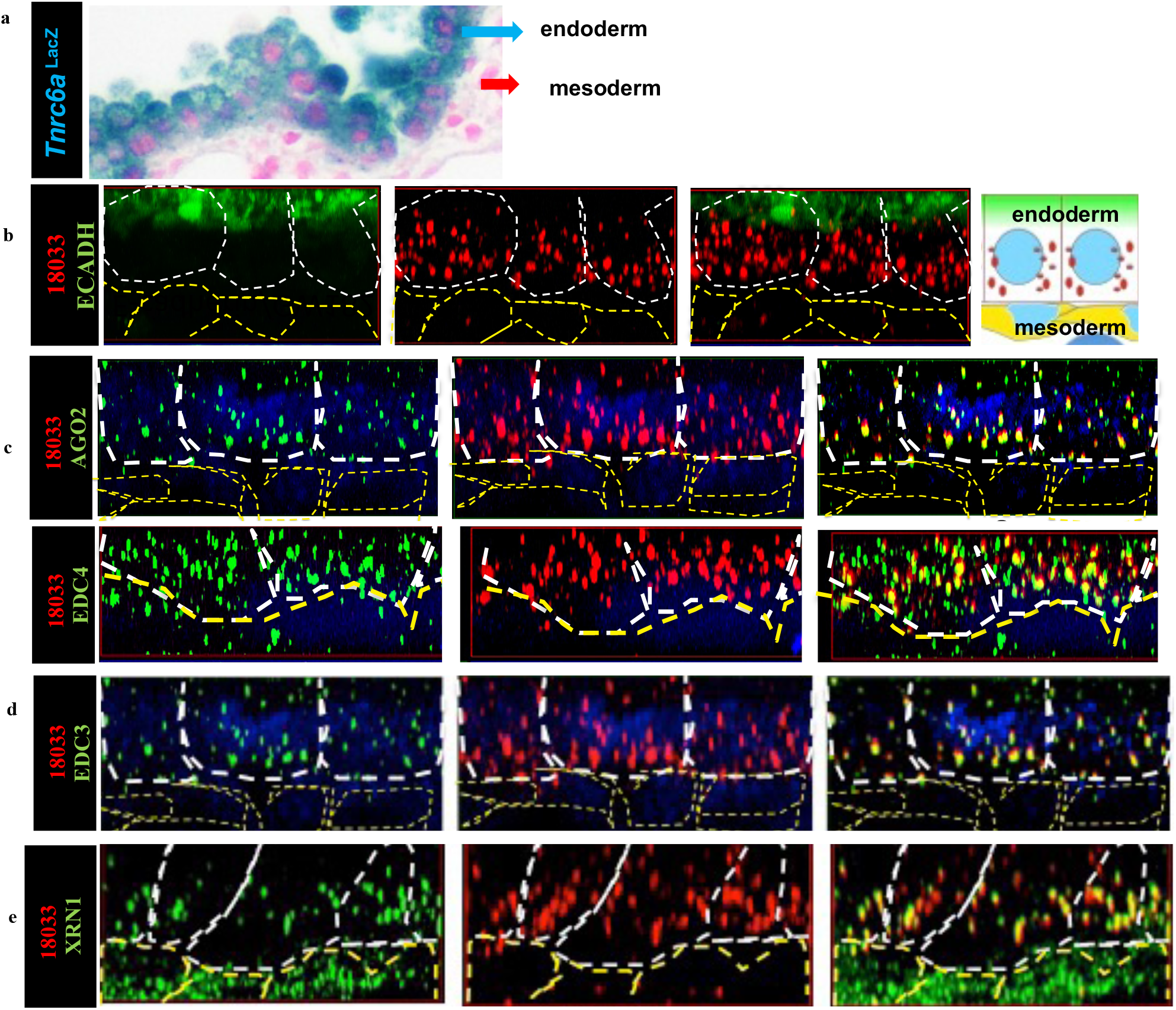
TNRC6A proteins concentrate in P-bodies containing AGO2 and enzymes for mRNA degradation in E9.5 mouse yolk sac. a) β-galactosidase staining of E9.5 yolk sac from Tnrc6a ^lacZ^ reporter mice showing selective expression in the endoderm. b-e) IF co-staining of TNRC6A (human index serum) with E-Cadherin, AGO2, EDC4, EDC3, XRN1 in yolk sacs. Cell boundaries are outlined by dashed lines (white: endoderm; yellow: mesoderm). Extensive overlap of TNR6A with AGO2 and mRNA degradation enzymes (EDC4, EDC3, XRN1) in P-bodies of the yolk sac endoderm (E-Cadherin positive). Y-Z maximum intensity projection confocal images. DAPI (blue) : nuclei.

**Supplementary Fig. 8.**
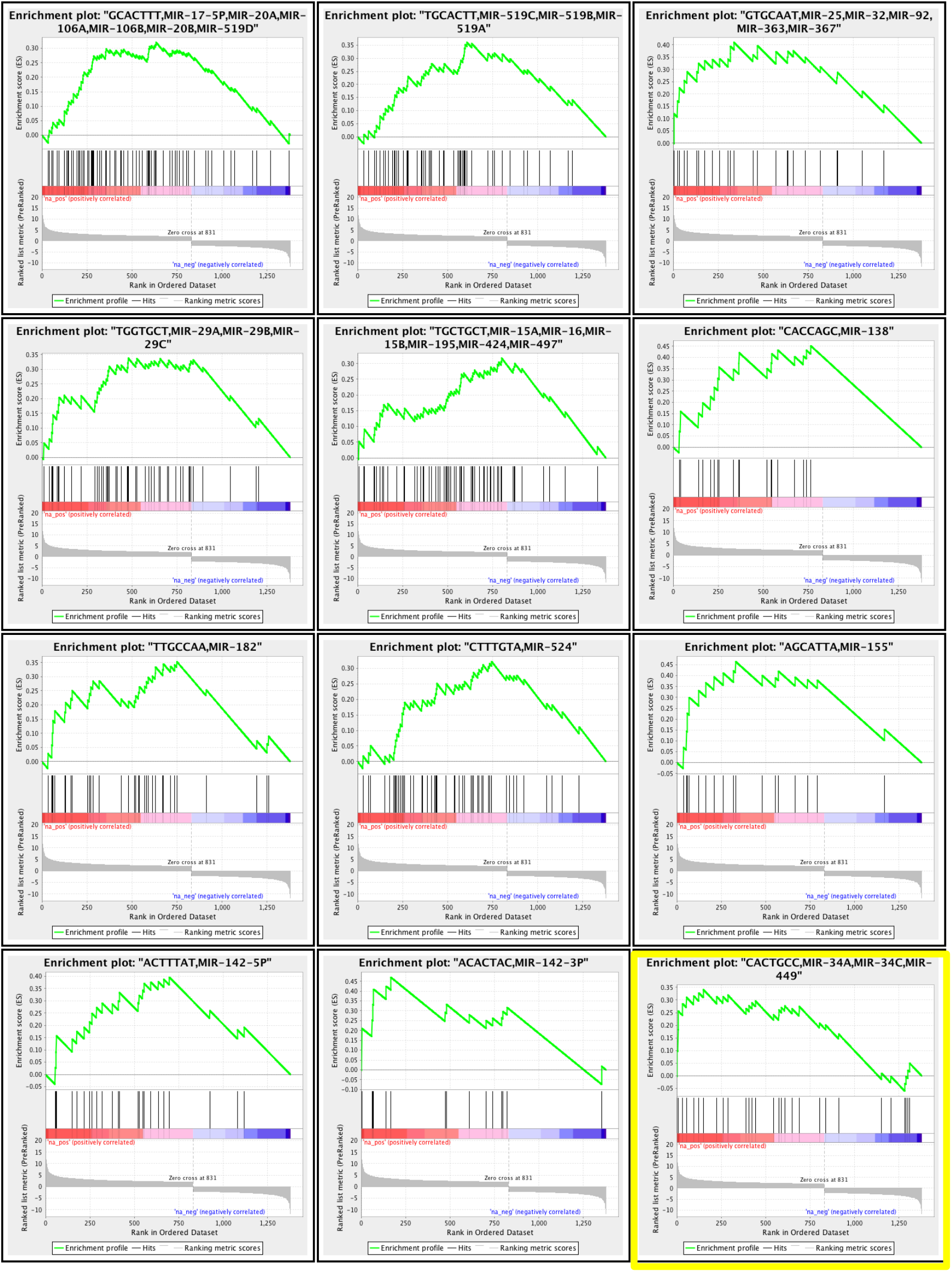
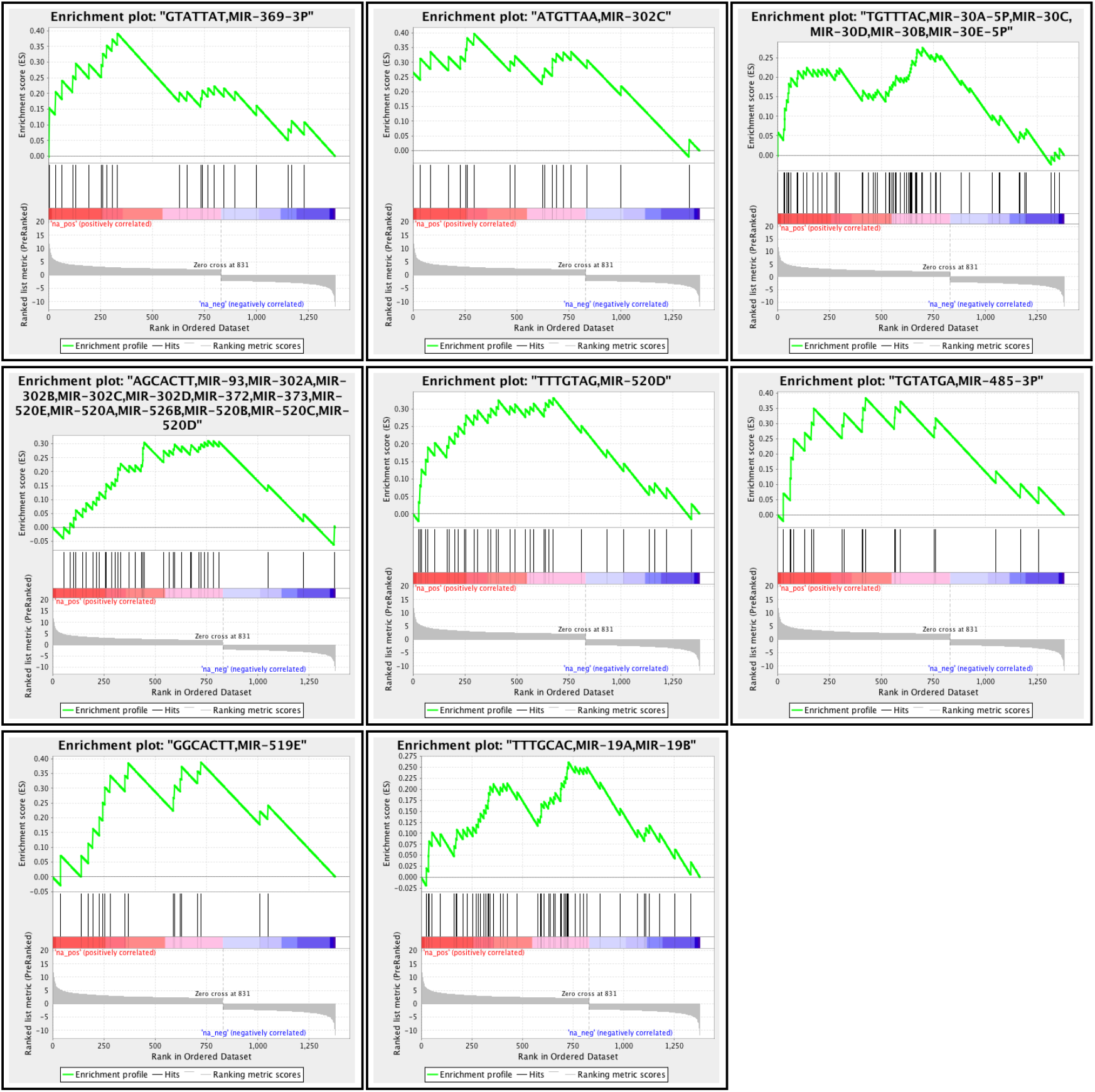
miRNA targets are enriched among genes upregulated in *Tnrc6a*^*null*^ yolk sac. miR34-449 highlighted

**Supplementary Fig. 9.**
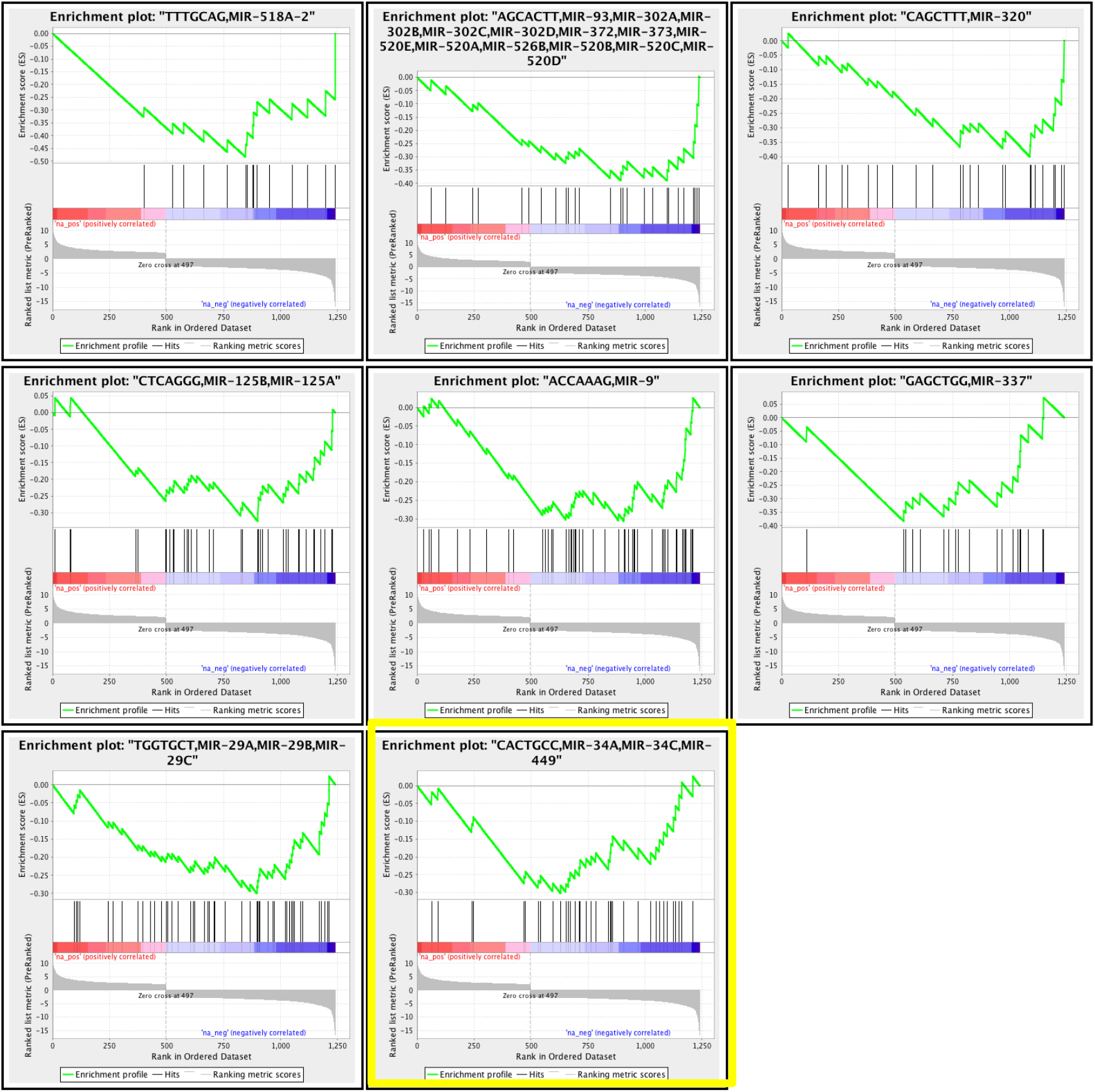
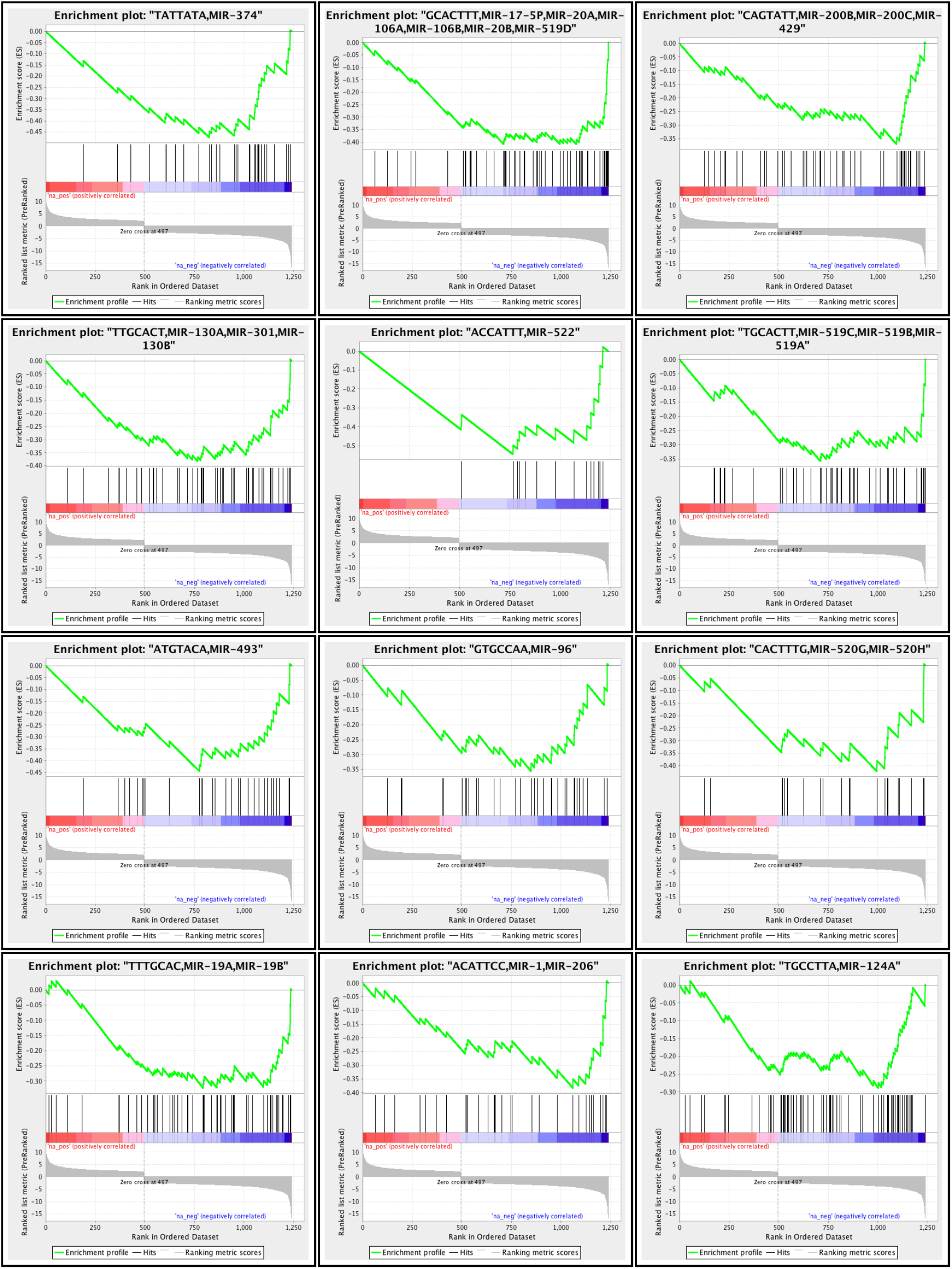
miRNA targets are enriched among genes downregulated in E18.5 *Tnrc6a*^*null*^ lungs. miR34-449 highlighted.

**Supplementary Fig. 10.**
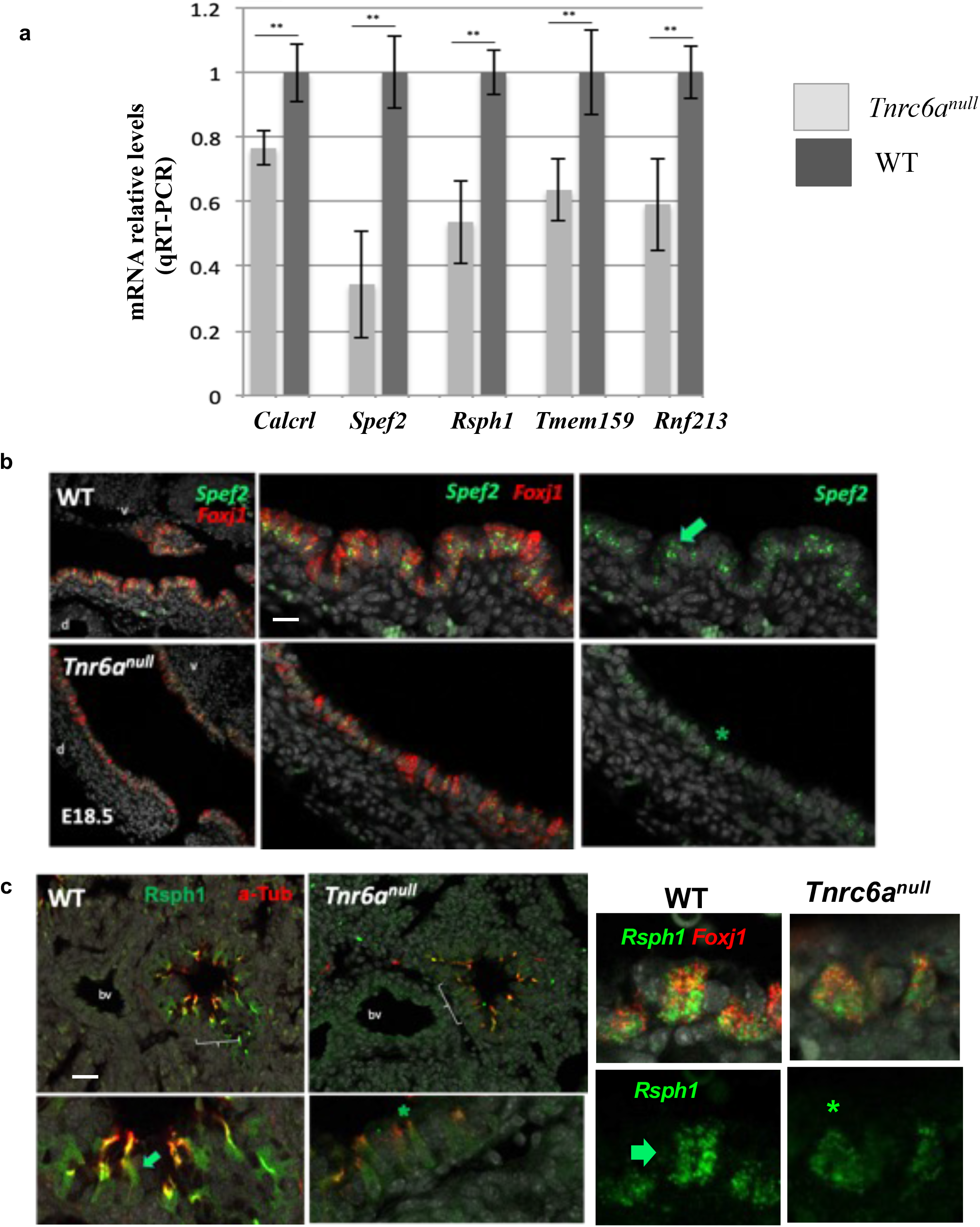
Decreased expression of miR-34/449 targets in Tnrc6a^null^ lungs. a) qRT-PCR confirmation of gene targets of miR-34/449 downregulated in *Tnrc6a*^null^ lungs. Bars are mean ± SEM (n=3 per group); Student’s t test’ ** p<0.01. (b) *Spef2* and *Foxj1* dual in situ hybridization (ISH, RNAscope) showing strong *Spef2* signals (arrow) in MCCs of WT, reduced in MCCs of *Tnrc6a*^null^ (representative large airway from both groups: d, dorsal; v, ventral). (c) Rsph1 and ac-a-Tubulin IF showing strong Rsph1 signals in WT MCCs (arrow bottom left) reduced in *Tnrc6a*^null^ airways (* bottom right). Right panel: Double ISH of *Rsph1-Foxj1* confirming strong Rsph1 signals in WT MCCs, decreased in mutant MCCs. (b, c) representative images of n=2 lungs per group. Bars in b, c = 12um and 15 um, respectively.

**Supplementary Fig. 11.**
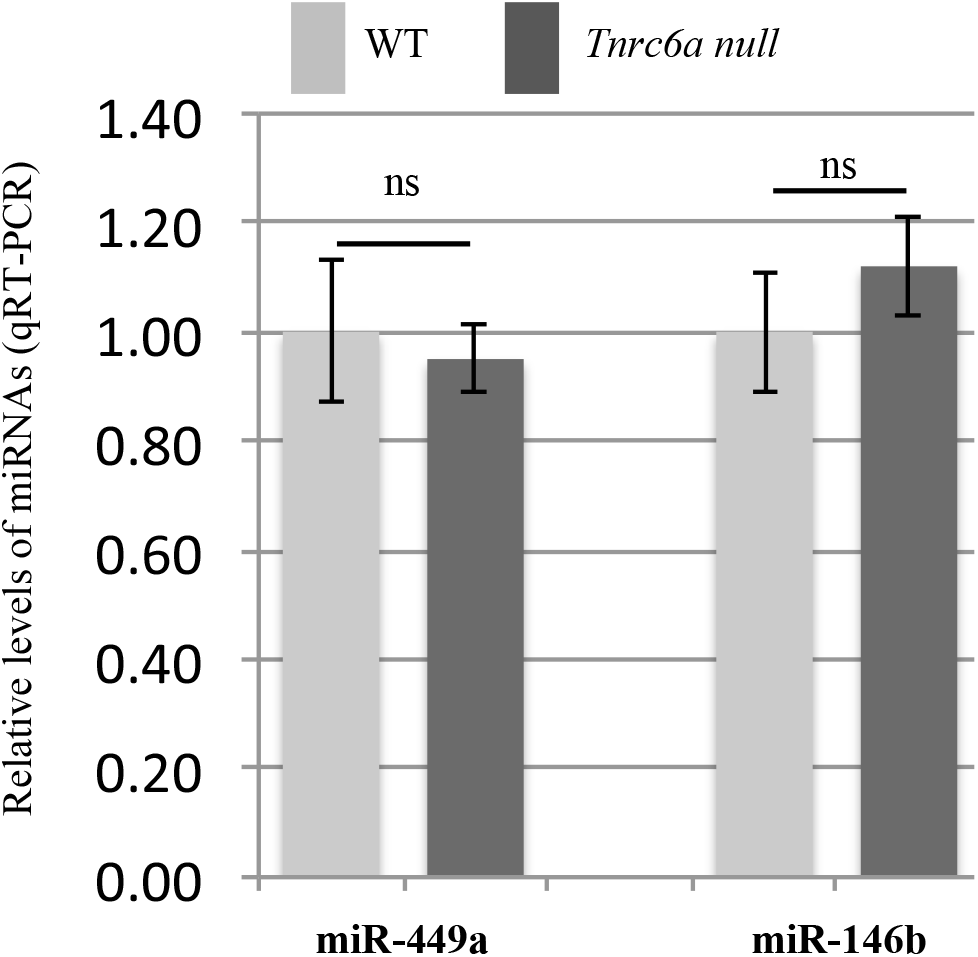
Levels of miR-449a and miR-146b are not altered in *Tnrc6a* ^*null*^ lungs. Bars are mean ± SD of values from n=3 lungs per group. Student’s t test (n=3/group). ns = non-significant.

**Supplementary Table 1.**
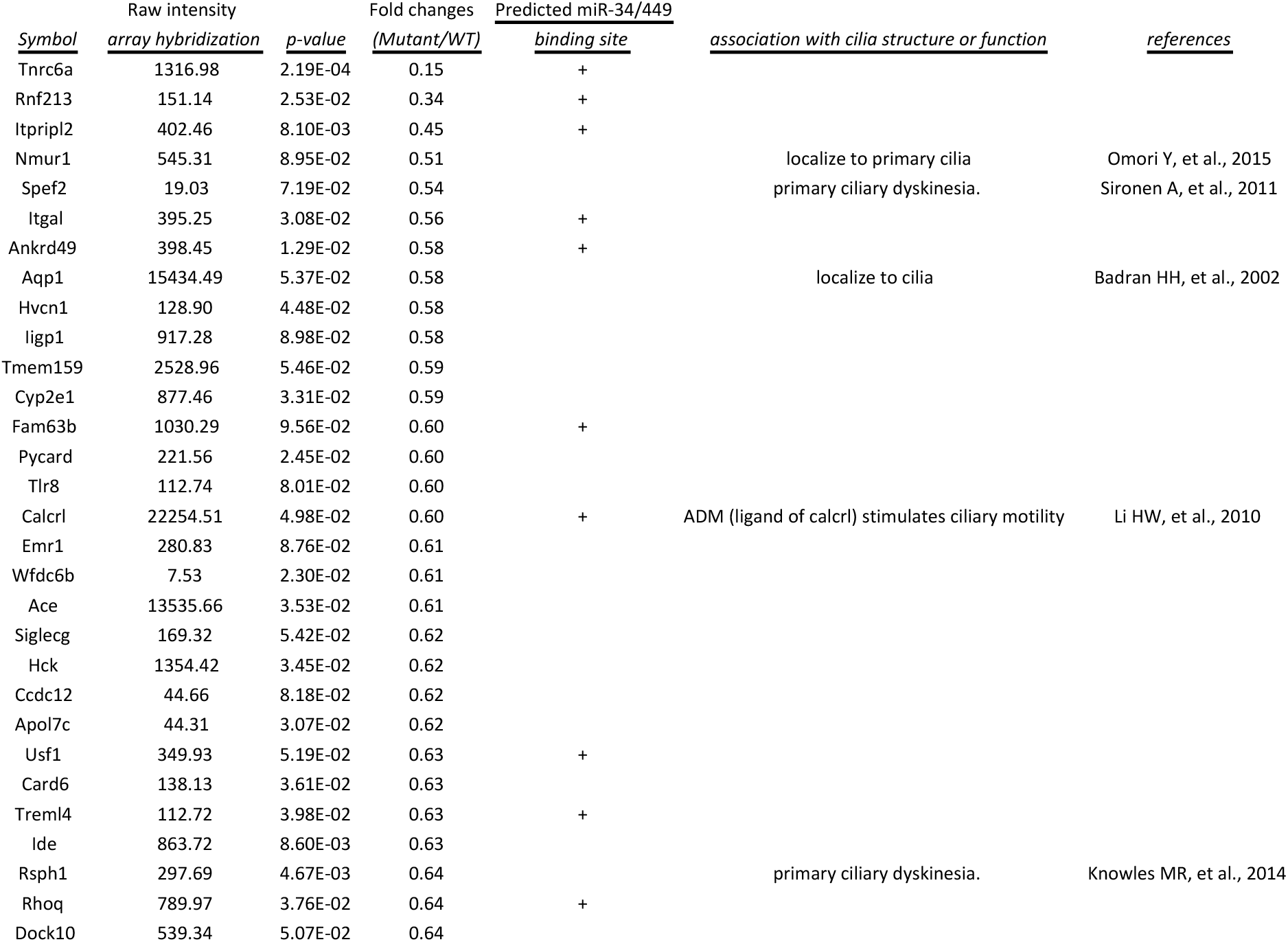
Genes most downregulated in *Tnrc6a* null lungs

